# New insights into the classification of the RAC1 P29S hotspot mutation in melanoma as an oncogene

**DOI:** 10.1101/2025.03.04.637030

**Authors:** Amin Mirzaiebadizi, Mohammad Reza Ahmadian

## Abstract

The RAC1^P29S^ hotspot mutation, prevalent in melanoma, drives tumorigenesis by enhancing molecular interactions and hyperactivating key signaling pathways, making it a compelling target for cancer therapy. This study provides a comprehensive biochemical characterization of RAC1^P29S^ compared to wild-type RAC1 and mutations T17N and F28L. The P29S mutation significantly impairs nucleotide binding to guanosine triphosphate (GTP) and guanosine diphosphate, accelerating intrinsic nucleotide exchange. While minimally affecting regulation by guanosine dissociation inhibitor 1, RAC1^P29S^ exhibits reduced activation via diffuse B-cell lymphoma family guanine nucleotide exchange factors but retains effective activation by dedicator of cytokinesis 2. Critically, the P29S mutation severely impairs GTPase-activating protein-stimulated GTP hydrolysis, most likely contributing to RAC1^P29S^ hyperactivation by prolonging its GTP-bound form. RAC1^P29S^ displays a stronger binding affinity for IQ motif-containing GTPase-activating protein 1 than for p21-activated kinase 1, highlighting the role of the former in scaffolding RAC1^P29S^-driven signaling. In serum-starved cells, RAC1^P29S^ predominantly adopts an active GTP-bound state. RAC1^P29S^ overexpression activates key cancer-associated pathways, including extracellular signal-regulated kinase and p38 mitogen-activated protein kinase, reinforcing its role as an oncogenic driver in melanoma. These insights suggest potential therapeutic targets for melanoma treatment, including RAC1 regulators and modulators.

**Graphical abstract:** 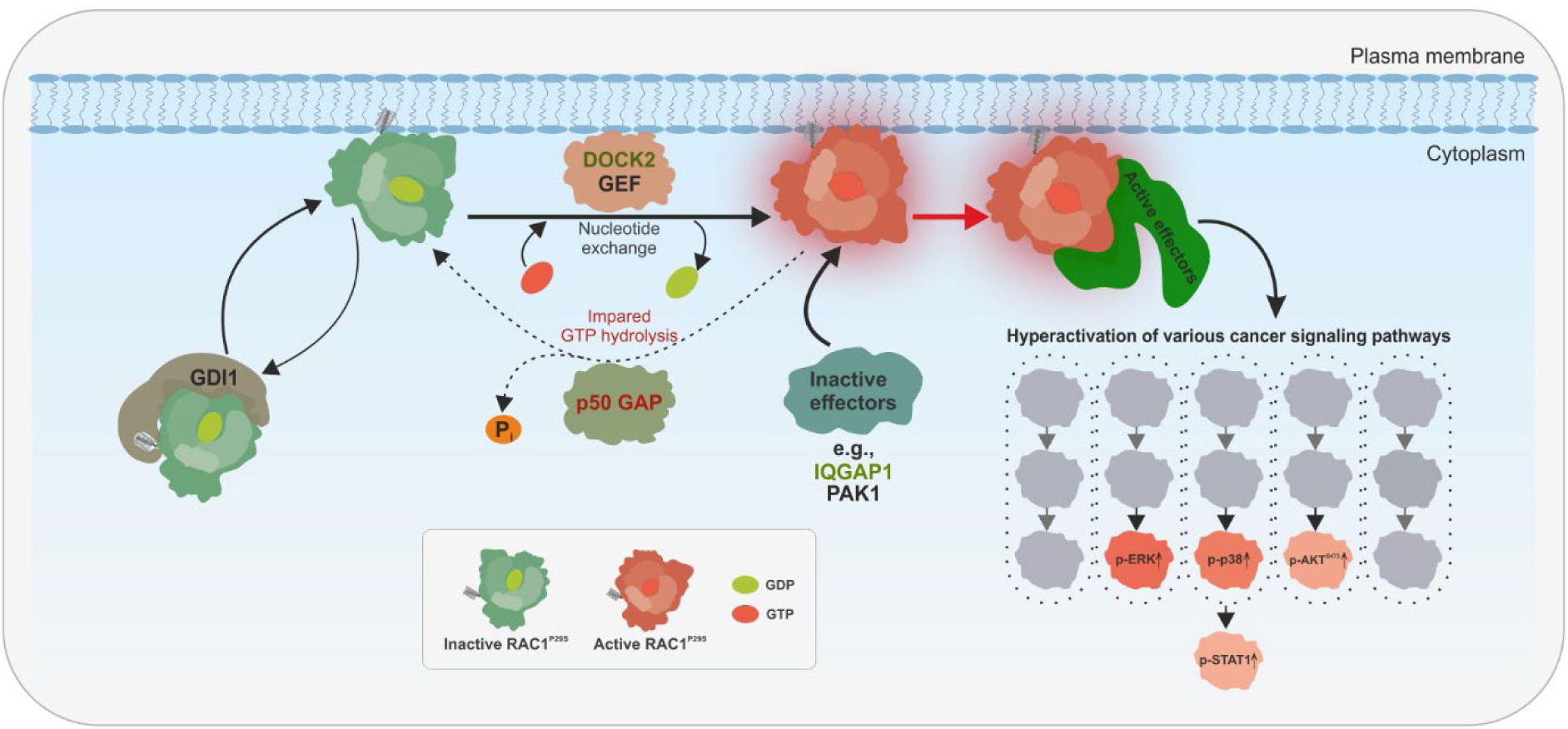

A model of RAC1^P29S^ activation and signaling in cancer cells. RAC1^P29S^ remains in an inactive GDP-bound state in the cytoplasm where GDI1 prevents its membrane association. Upon stimulation, GEFs, primarily DOCK2, activate RAC1^P29S^ by promoting GDP-GTP exchange, facilitating its transition to the active GTP-bound state and initiating downstream signaling. RAC1^P29S^ binds preferentially to IQGAP1 over PAK1, reflecting a shift in effector interactions. IQGAP1 acts as a scaffolding protein, spatially modulating RAC1^P29S^-driven signaling and amplifying its effects. Under normal conditions, GAPs such as p50GAP regulate RAC1 by accelerating GTP hydrolysis, thereby maintaining its dynamic activation cycle. However, the P29S mutation severely impairs p50GAP-mediated hydrolysis, leading to accumulation of RAC1^P29S^ in its GTP-bound state and loss of temporal regulation. This persistent activation hyperactivates downstream effectors and promotes cancer-associated pathways, including ERK and p38 MAPK, which drive cell growth, survival, invasion and metastasis.

## Introduction

As a key member of the RHO guanosine triphosphatase (GTPase) family, RAC1 functions as a molecular switch, cycling between an inactive guanosine diphosphate (GDP)-bound form and an active guanosine triphosphate (GTP)-bound form (1). This switch relies on two essential processes—GDP/GTP exchange and GTP hydrolysis—which induce structural changes in the switch I (amino acids 29–42) and switch II (amino acids 62–68) regions (2). These functions are regulated by guanine nucleotide exchange factors (GEFs) and GTPase-activating proteins (GAPs) (3–5). The RHO GEF family includes the structurally distinct dedicator of cytokinesis (DOCK) and diffuse B-cell lymphoma (DBL) subfamilies (1, 6, 7). In addition, guanine nucleotide dissociation inhibitors (GDIs) selectively bind geranylgeranylated RAC1, controlling its membrane localization (8).

RAC1 and its isoform RAC1B (9) and paralogs RAC2 and RAC3 (10) activate diverse signaling pathways through direct interaction with effector proteins (1). These interactions regulate essential cellular processes, including motility, oxidative stress, and inflammation (11). GTP-bound RAC1 binds effectors, activating kinases like p21-activated kinase 1 (PAK1) and scaffolding proteins like IQ motif-containing GTPase-activating protein 1 (IQGAP1) (1). Dysregulation (12, 13) or gain-of-function mutations in RAC genes (14, 15) can hyperactivate RAC signaling, altering cellular responses and contributing to cancer. This dysregulation contributes to various pathological conditions, including cancer (16), and other pathological conditions, including metabolic, neurodegenerative, cardiovascular, inflammatory, and infectious diseases (11).

The proline 29 to serine (P29S) mutation in RAC1 is the third most common hotspot mutation in melanoma, following BRAF V600E and NRAS Q61R (17). Despite its prevalence, the regulatory functions driving RAC1^P29S^ pro-tumorigenic effects remain poorly understood (18). Functional studies show that RAC1^P29S^ enhances effector binding, including PAK1 and mixed lineage kinase 3 (MLK3), promoting melanocyte proliferation and migration (19, 20). Additionally, RAC1^P29S^ inhibits invadopodia function (21), abolishes haptotaxis (22), drives dedifferentiation in melanoma, contributes to BRAF inhibitor resistance (23–25), and facilitates immune evasion via programmed death-ligand 1 (PD-L1) upregulation through the RAC1^P29S^-PAK1 axis (17). This immune evasion is mediated by the RAC1^P29S^-PAK1 axis, which promotes the G2/M cell cycle transition through phosphorylation of Aurora kinase A and polo-like kinase 1 (PLK1) (26) and inactivates neurofibromin 2 (NF2)/Merlin, promoting proliferation, metastasis, and drug resistance (27). Furthermore, while BRAF^V600E^ suppresses cell migration, extracellular signal-regulated kinase (ERK) pathway inhibition accelerates migration and invasion in BRAF^V600E^- and mutant RAS-driven tumors (28). Although RAC1 is a critical therapeutic target in melanoma, its undruggable nature poses a significant challenge for targeting RAC1^P29S^ (11, 29–35).

Initial studies using radiolabeled nucleotide filter binding assays or thin-layer chromatography compared the basal GDP/GTP exchange and GTP hydrolysis of RAC1^P29S^ with RAC1^WT^. Davis et al. reported increased GTP dissociation for RAC1^P29S^ (36), while Kawazu et al. observed increased GDP dissociation but not GTP dissociation (37). Both studies concluded that GTP hydrolysis remained unchanged. However, these and other overexpression studies alone cannot classify RAC1^P29S^ as spontaneously activating, self-activating, fast cycling, constitutively active, or oncogenic (Box 1) (20, 21, 36, 38, 39). Some of these classifications are derived from assumptions about the phenylalanine 28 to leucine (F28L) mutant of RAC1. Although RAC1^F28L^ is not extensively studied, it is described as a fast-cycling mutant, analogous to CDC42^F28L^, capable of spontaneous nucleotide exchange without GEF activation while retaining full GTPase activity (40). Another widely studied mutant, threonine 17 to aspargine (T17N), is a dominant negative mutant with T17 in the phosphate-binding loop (P-loop), a region critical for nucleotide binding, while F28 and P29 reside at the N-terminus of switch I. The P-loop and switch I are essential for RAC1 nucleotide binding and hydrolysis (1). Biophysical and biochemical studies, supported by molecular dynamics simulations, indicate that the P29S mutation increases switch I flexibility, adopting an open conformation that facilitates rapid GDP/GTP exchange in RAC1 (20, 36, 41, 42).

This study comprehensively characterizes RAC1^P29S^ at three levels: intrinsic properties, regulation, and effector interaction. At the intrinsic level, we analyzed its nucleotide exchange, GTP hydrolysis, and binding affinities for GDP and GTP. Studies on regulatory mechanisms examined its interaction with DBL and DOCK family GEFs, p50 Rho GTPase-activating protein (p50GAP)-mediated GTP hydrolysis, and GDI1-mediated regulation. Effector interactions focused on key proteins, including PAK1, a kinase, and IQGAP1, a scaffolding protein modulating RAC1 signaling. Active GTPase pull-down experiments under serum-stimulated and serum-starved conditions provided further insights into the GTP-bound state of RAC1^P29S^ in cells. Comparative analyses with RAC1^WT^, RAC1^T17N^, and RAC1^F28L^ revealed distinct properties of RAC1^P29S^, including a fast intrinsic nucleotide exchange rate, DOCK2-mediated exchange facilitation, and severely impaired p50GAP-stimulated GTP hydrolysis. These biochemical and functional differences result in hyperactivation and preferential signaling through IQGAP1 rather than PAK1. Taken together, these findings highlight potential therapeutic targets in melanoma, including DOCK2, p50GAP, and IQGAP1.

## Material and Methods

### Constructs

Human RAC1 wild-type (RAC1^WT^; accession no. P63000) and its mutants T17N, F28L, and P29S were expressed as N-terminal glutathione S-transferase (GST)-tagged fusion proteins using pGEX vectors (pGEX-2T and pGEX-4T-1). The same system was used to express regulators and effectors, including full-length GDI1, the Dbl homology-pleckstrin homology (DH-PH) tandem domains of T-lymphoma invasion and metastasis-inducing protein 1 (TIAM1), vav guanine nucleotide exchange factor 2 (VAV2), son of sevenless homolog 1 (SOS1), and phosphatidylinositol-3,4,5-trisphosphate-dependent Rac exchanger 1 (PREX1); the GAP domain of p50GAP; the C-terminal 794-amino acid region of IQGAP1; and the RAC1 binding domain (RBD) of PAK1. Additional constructs included His-tagged IQGAP1 (pET-23b+ vector) and His6-small ubiquitin-like modifier (SUMO)-tagged DOCK2 Dock homology region 2 (DHR2) domain (pOPINS vector). RAC1 constructs with N-terminal tandem decahistidine triple-flag tags were cloned into the pcDNA-3.1 vector for eukaryotic expression. Detailed deconstructs description including accession numbers and amino acid sequences, are available in the Supplementary information.

### Proteins

All proteins were purified as described previously (3, 5, 9, 10, 43). Briefly, Escherichia coli strains were transformed for protein expression, lysed, and subjected to affinity purification using GST or His tags. GST tags were cleaved when necessary, and proteins were buffer-exchanged into optimized storage buffers. Purity was confirmed by SDS-PAGE and Coomassie staining (Supplementary Fig. S1), which shows the purified proteins used in this study. Proteins were stored at -80 °C. Detailed procedures are available in the Supplementary Materials.

### Preparation of nucleotide-free and fluorescent nucleotide-bound GTPases

As previously described, nucleotide-free GTPases were prepared through sequential treatment with alkaline phosphatase and snake venom phosphodiesterase (44, 45). Fluorescent GDP- and GppNHp-bound GTPases were generated by incubating nucleotide-free proteins with mant-labeled nucleotides (mdGDP and mGppNHp). Protein concentrations were quantified by HPLC using NAP-5 buffer exchange columns. Samples were stored at −80 °C. Detailed procedures are provided in the Supplementary Materials.

### Fluorescence kinetic measurements

Fluorescence-based kinetic measurements for long-term and rapid reactions were performed using a Horiba Fluoromax-4 fluorimeter and a stopped-flow spectrophotometer (Applied Photophysics SX20), as described (43–46). Excitation and emission wavelengths were set according to the fluorophore-specific properties of mant- and tamra-labeled nucleotides. Detailed experimental conditions are provided in the Supplementary Materials.

### Nucleotide-binding assay

The nucleotide-binding properties of RAC1 GTPases were assessed by stopped-flow fluorimetry, as described (47). Nucleotide association and dissociation rates were measured using fluorescent nucleotides (mdGDP and mGppNHp) and varying RAC1 concentrations. Association (k_on_) and dissociation (k_off_) rate constants were determined, and equilibrium dissociation constants (K_d_) were calculated as described in Box 2. Detailed procedures are provided in the Supplementary Materials.

### GEF-catalyzed nucleotide dissociation assay

The GEF-catalyzed nucleotide exchange reaction was monitored by stopped-flow fluorimetry, as described (45). Reactions were performed with mGDP-bound RAC1 and excess non-fluorescent nucleotide in the presence of GEFs from the DBL and DOCK families. Observed rate constants were analyzed using a single-exponential model in Origin software. Detailed procedures are provided in the Supplementary Materials.

### Intrinsic and GAP-stimulated GTP-hydrolysis assays

The intrinsic GTP hydrolysis rate of RAC1 proteins was determined by high-performance liquid chromatography (HPLC), as described (45). Reactions were performed with nucleotide-free RAC1 and GTP in buffer C at 25 °C, and catalytic rate constants (k_cat_) were calculated using Origin software. GAP-stimulated hydrolysis rates were measured by stopped-flow fluorimetry using tamra-GTP, as described (48). Detailed procedures are provided in the Supplementary Materials.

### Protein-protein interaction kinetics

The interaction of RAC1 with GST-GDI1, GST-PAK1 RBD, and His-IQGAP1 C794 was analyzed by stopped-flow fluorimetry to determine k_on_, k_off_, and K_d_ values, as described (43). Binding assays were performed using mdGDP- and mGppNHp-bound RAC1 with varying protein concentrations, and rate constants were calculated using linear regression and single-exponential fits. Detailed procedures are provided in the Supplementary Materials.

### Fluorescence polarization

Fluorescence polarization was used to determine the binding affinity between RAC1 and effector proteins, as described (43). Assays were performed with mGppNHp-bound RAC1 (1 μM) and titrated effectors in buffer containing 30 mM Tris-HCl (pH 7.5), 50 mM NaCl, 5 mM MgCl_2_, and 3 mM DTT at 25 °C. K_d_ values were calculated by fitting binding curves to a quadratic ligand binding equation. Detailed procedures are provided in the Supplementary Materials.

### Cell culture and transfection

HEK-293T cells were cultured under serum-stimulated and serum-starved conditions in DMEM supplemented with 10% FBS and 1% penicillin/streptomycin. RAC1 constructs with N-terminal 10 His-triple flag tags were transfected using TurboFect™ (Thermo Fisher Scientific) according to the manufacturer’s protocol. Cells were harvested and lysed, and protein concentrations were measured using the Bradford assay. Detailed protocols, including buffer compositions, are provided in the supplementary information.

### *In vitro* pull-down assays

Pull-down assays were conducted to assess RAC1 binding to PAK1 RBD and IQGAP1 C794. GST-PAK1 RBD and His-IQGAP1 C794 were immobilized on glutathione-agarose and His-Mag Sepharose Ni beads, respectively. RAC1 proteins were incubated with the beads, washed, eluted, and analyzed by SDS-PAGE followed by immunoblotting. Detailed protocols are provided in the Supplementary Materials.

### Active GTPase pull-down assay

A pull-down assay was performed to assess GTP-bound active RAC1 levels in transiently transfected HEK-293T cells under serum-stimulated and serum-starved conditions, as described (49). GST-PAK1 RBD- and GST-IQGAP1 C794-coupled beads were prepared, incubated with lysates, washed, and analyzed by SDS-PAGE and immunoblotting. Detailed protocols are provided in the Supplementary Materials.

### Antibodies and Immunoblotting

Primary and secondary antibodies were diluted in TBST with a blocking buffer. The antibodies included α-RAC1, α-6x-His, α-flag, α-γ-tubulin, α-p-ERK1/2, α-t-ERK1/2, α-p-AKT, α-t-AKT, α-p-p38 MAPK, α-p38 MAPK, α-p-STAT1, α-STAT1, α-GAPDH, and α-GST. Immunoblots were visualized using the Odyssey® XF Imaging System. Detailed antibody lists and protocols are provided in the Supplementary Materials.

### Statistical Analysis

Data in bar graphs represent mean ± S.D., with replicate numbers detailed in figure legends. Immunoblot intensities were quantified using Image Studio Lite 5.2. For pull-down assays, data were normalized based on RAC1-to-effector ratios, and active RAC1 levels were calculated as described in Fig. 4B. Downstream signaling data were normalized to phosphorylated-to-total protein ratios and adjusted to GAPDH levels as a loading control. Statistical significance was determined using one-way ANOVA followed by Tukey’s test (*P ≤ 0.05, **P ≤ 0.01, ***P ≤ 0.001, ****P ≤ 0.0001). Detailed normalization methods and calculations are provided in the Supplementary Materials.

## Results

### P29S significantly impairs the nucleotide binding of RAC1

Two sets of real-time kinetic measurements were performed to investigate the impact of the P29S mutation on nucleotide binding affinity. The first measured the association of mdGDP and mGppNHp with nucleotide-free (n.f.) RAC1 (Fig. 1A), while the second analyzed the dissociation of these nucleotides from RAC1 (Fig. 1B). The fluorescent analog mdGDP was used as a substitute for GDP, and the non-hydrolyzable mGppNHp replaced GTP. The RAC1 variants included WT, T17N, F28L, and P29S.

**Figure 1.**
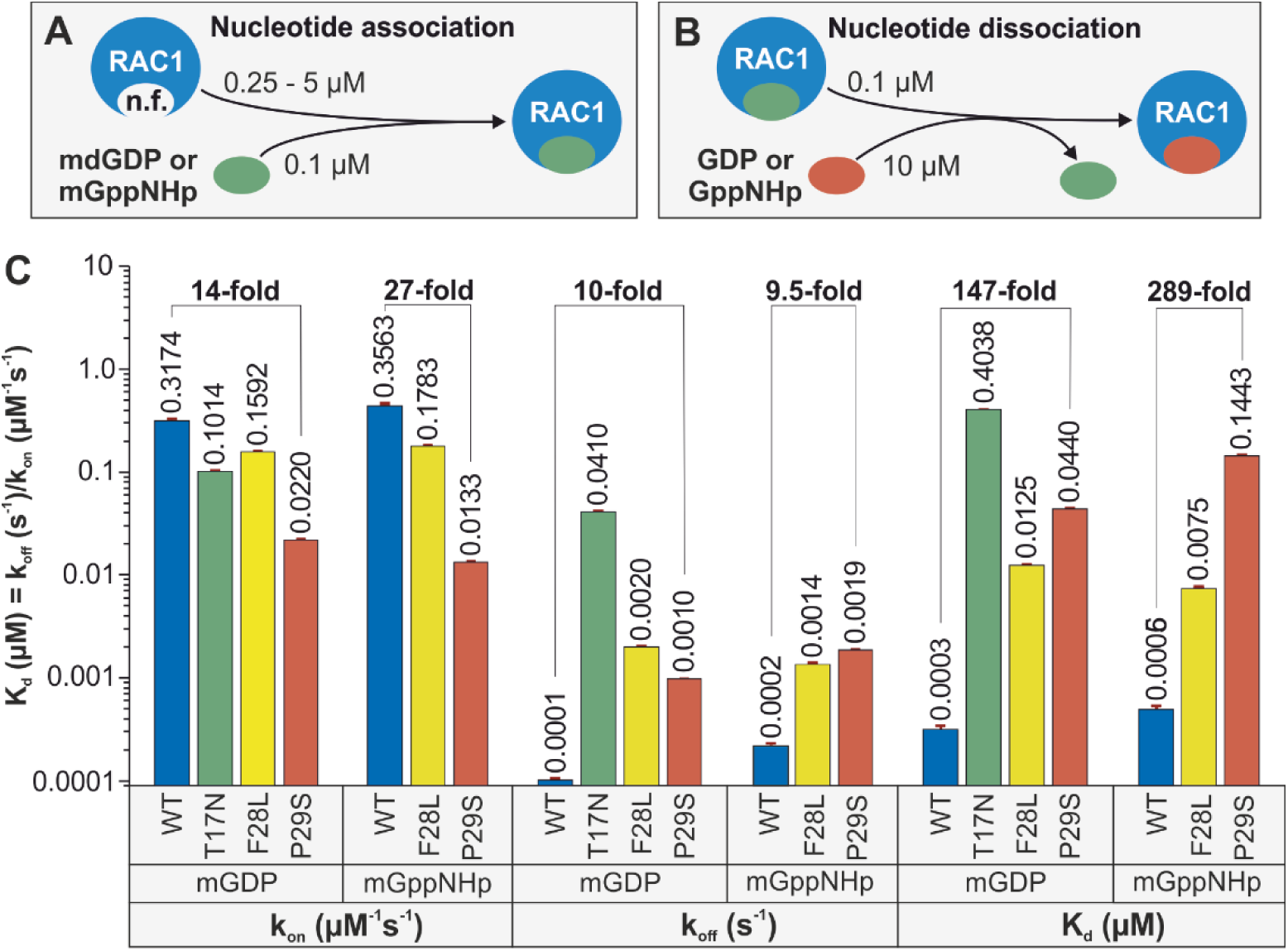
Severe impairment of the GDP/GTP binding properties of RAC1^P29S^. The kinetics of association (**A**) and dissociation (**B**) of fluorescent mdGDP and mGppNHp with RAC1 proteins were measured as illustrated. (**C**) Kinetic rate constants for association (k_on_) and dissociation (k_off_), as well as the dissociation constant (K_d_), calculated from the k_off_/k_on_ ratio, reveal substantial effects of the P29S mutation on the binding of mdGDP and mGppNHp to RAC1. These effects differ markedly from those observed for the T17N and F28L substitutions. This impaired binding may contribute to the accelerated intrinsic nucleotide exchange observed in RAC1^P29S^. All k_on_, k_off_, and K_d_ values, presented as bar graphs, represent the average of three to six measurements and are reported as means ± SD.

Binding of nucleotides to n.f. RAC1 induced a rapid fluorescence increase, with k_obs_ values rising proportionally with n.f. RAC1 concentrations (supplementary Fig. S2 and S3, left panels), which depict the interaction of mdGDP and mGppNHp with RAC1 at increasing concentrations. The k_on_ values for mdGDP and mGppNHp binding were derived from linear fits of kobs values across protein concentrations (supplementary Fig. S2 and S3, middle panels), where k_on was determined by plotting observed rate constants from exponential fits of association data against the corresponding RAC1 concentrations. A bar graph of kon values showed significant differences in nucleotide association among RAC1 variants (Fig. 1C). The P29S mutation notably reduced the association of mdGDP and mGppNHp with RAC1 by 14-fold and 27-fold, respectively, compared to RAC1^WT^.

A decrease in fluorescence was observed during nucleotide dissociation from RAC1 proteins in the presence of excess free GDP (supplementary Fig. S2 and S3, right panels), which depict the dissociation kinetics of mdGDP and mGppNHp from RAC1 proteins. The k_off_ values, derived from single exponential fits of the dissociation data, are shown as bar graphs (Fig. 1C). RAC1^P29S^ and RAC1^F28L^ exhibited intrinsic nucleotide dissociation rates 10- and 20-fold faster than RAC1^WT^, respectively. RAC1^T17N^ showed the fastest mdGDP dissociation rate, 410-fold higher than RAC1^WT^, resulting in a significantly reduced K_d_ and a 1346-fold decrease in binding affinity, highlighting its dominant-negative effect (see Box1 for Definitions). Additionally, due to extremely rapid association and dissociation rates, mGppNHp kinetics for RAC1^T17N^ could not be determined using stopped-flow fluorimetry (supplementary Fig. S3, lower panel), which presents fluorescence spectrophotometry-based measurements confirming its rapid nucleotide exchange properties.

Nucleotide-binding affinity (K_d_) was calculated using kinetic parameters for dissociation and association reactions. RAC1^WT^ displayed tight binding affinities for mdGDP and mGppNHp, with Kd values of 0.3 nM and 0.6 nM, respectively. These affinities were significantly reduced for RAC1^T17N^, followed by RAC1^P29S^ and RAC1^F28L^ (Fig. 1C). Due to rapid kinetics, the mGppNHp binding affinity for RAC1^T17N^ could not be determined (supplementary Fig. S3, lower panel), where fluorescence measurements demonstrated its inability to be analyzed via standard stopped-flow techniques. RAC1^P29S^ showed markedly impaired nucleotide binding, with 147-fold and 289-fold lower affinities for mdGDP and mGppNHp, respectively. These findings suggest that RAC1^P29S^’s impaired binding properties likely drive its accelerated intrinsic nucleotide exchange, although further structural studies on its interactions with regulators and effectors are needed to elucidate its aberrant behavior.

### Only the T17N mutation significantly impairs GDI1 activity

We recently developed a fluorescence-based method to monitor RAC1-GDI1 interactions (8). Our results showed that GDI1 binding, essential for GDI-mediated membrane translocation, does not differentiate between non-prenylated and prenylated RAC1. Real-time kinetic measurements evaluated the association and dissociation kinetics of GDI1 with mdGDP-bound RAC1 (Fig. 2A, left panel; supplementary Fig. S4), which presents the binding of RAC1 to GST-GDI1 across increasing concentrations, followed by kinetic analysis. Corresponding rate constants are shown in Figure 2A, right panel. RAC1^F28L^ and RAC1^P29S^ exhibited k_on_ and k_off_ values comparable to RAC1^WT^, with slightly reduced GDI1 binding affinity for RAC1^P29S^. In contrast, RAC1^T17N^ displayed a 439-fold decrease in GDI association and an 18-fold reduction in dissociation, leading to a significantly decreased binding affinity compared to RAC1^WT^.

**Figure 2.**
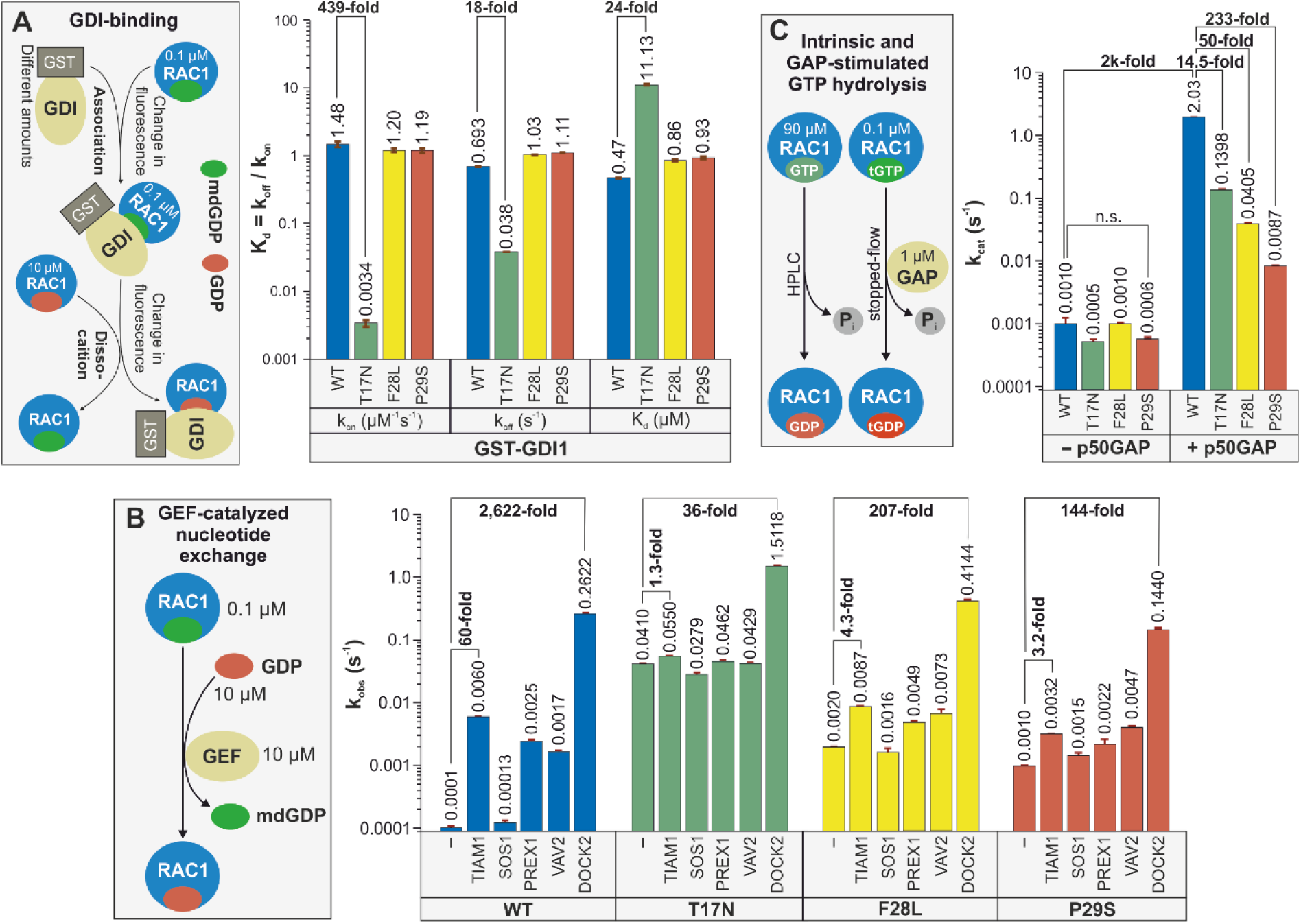
Effects of mutations on the regulation of RAC1 by GDI1, various GEFs, and p50GAP. (**A**) Minimal effect of the P29S mutation on the RAC1-GDI1 interaction. The principle behind the kinetic measurements of the association of GST-GDI1 with RAC1 proteins and its dissociation is illustrated using a stopped-flow instrument. In these experiments, 0.1 µM mdGDP-bound RAC1 was rapidly mixed with increasing concentrations of GST-GDI1 to monitor the association kinetics. Dissociation kinetics were measured by rapidly mixing a complex of RAC1•mdGDP•GST-GDI1 with excess GDP-bound RAC1. Bar graphs from the stopped-flow analysis depict the association rates (k_on_) and dissociation rates (k_off_) of the GDI1 interaction from/with RAC1 proteins, as well as the dissociation constant (K_d_), calculated from the k_off_/k_on_ ratio. The analysis revealed a substantial reduction in GDI1 binding affinity for RAC1^T17N^ and a slight reduction for RAC1^P29S^. All kinetic data are based on the average of three to six measurements and are presented as mean ± SD. (**B**) Impairment of the catalyzed nucleotide exchange of RAC1^P29S^ by DBL proteins but not by DOCK2. The mdGDP-to-GDP exchange of RAC1 proteins was measured in the absence and presence of the DH-PH tandem of various DBL family members (TIAM1, SOS1, PREX1, and VAV2) and the DHR2 domain of DOCK2, a member of the DOCK family. The observed rate constants (k_obs_), shown as bar graphs, represent the average of three to six measurements and are displayed as means ± SD. (**C**) Severely impaired GAP-stimulated GTP hydrolysis reaction of RAC1^P29S^. The basal and p50GAP-stimulated GTP hydrolysis reactions were measured using HPLC and stopped-flow instruments, respectively. The determined catalytic rate constants (k_cat_), presented as bar graphs, are based on duplicate measurements for HPLC data and three to six measurements for stopped-flow data and are reported as means ± SD.

### RAC1^P29S^ is mainly activated by DOCK2 and not by DBL family GEFs

To assess GEF-mediated nucleotide exchange, we evaluated mdGDP dissociation from RAC1^WT^, RAC1^T17N^, RAC1^F28L^, and RAC1^P29S^ in the presence of various RAC1-selective GEFs, including DBL family members (TIAM1, VAV2, SOS1, PREX1) (3, 4) and DOCK family member DOCK2 (7, 50) (Fig. 2B, left panel; supplementary Fig. S5), which presents kinetic measurements of GEF-catalyzed mdGDP dissociation from RAC1 proteins. Fluorescence decay curves were fitted to a single exponential function to determine k_off_ values in the presence of each GEF. Substantial GEF activity against RAC1^WT^ was observed for the DH-PH domains of TIAM1, PREX1, and VAV2, but not SOS1, consistent with prior reports (3). This lack of SOS1 activity extended to RAC1 mutants. Our findings indicate that RAC1^P29S^ has slow basal nucleotide exchange with DBL proteins and is primarily activated by DOCK2. The DHR2 domain of DOCK2 exhibited 40-fold greater activity than TIAM1 against RAC1^WT^ and showed significant GEF activity for RAC1^P29S^ and other mutants (Fig. 2B). This evidence positions DOCK2 as the primary potential activator of RAC1^P29S^ in cancer cells, particularly in melanoma.

### The P29S mutation significantly impairs the GAP activity

GTP hydrolysis was evaluated using HPLC for intrinsic hydrolysis and stopped-flow fluorimetry for GAP-stimulated hydrolysis (Fig. 2C; supplementary Fig. S6), which presents measurements of both basal and GAP-stimulated GTP hydrolysis of RAC1 proteins. Intrinsic hydrolysis was assessed by quantifying relative GTP content via HPLC, while real-time hydrolysis in the presence of p50GAP was analyzed using stopped-flow fluorescence. RAC1^WT^ exhibited slow intrinsic GTP hydrolysis (k_cat_ = 0.001 s⁻¹), consistent across RAC1 mutants, including RAC1^P29S^ (Fig. 2C). In contrast, RAC1^P29S^ showed a dramatic reduction in GAP-stimulated hydrolysis, with k_cat_ dropping from 2.03 s⁻¹ for RAC1^WT^ to 0.0087 s⁻¹, a 233-fold decrease (Fig. 2C, right panel). T17N and F28L mutations also reduced GAP activity but to a lesser extent (14.5-fold and 50-fold, respectively). These results underscore the critical role of GAP in the temporal regulation of RAC1 activity, with diminished p50GAP activity prolonging RAC1^P29S^’s GTP-bound state and enhancing its signaling capacity.

### RAC1^P29S^ shows a significantly stronger binding affinity to IQGAP1 compared to PAK1

The diverse signaling activities of RAC1 in human cells and cancers are primarily mediated through its interactions with downstream effectors. To evaluate the impact of the P29S mutation on effector binding under cell-free conditions, we examined its interaction with two well-characterized RAC1 effectors: the RAC1 binding domain (RBD) of the serine/threonine kinase PAK1, a key downstream kinase, and the C-terminal 794 amino acids (C794) of the scaffolding protein IQGAP1, a critical accessory protein (9, 10, 51, 52).

The binding properties of RAC1 mutants to PAK1 RBD were assessed using a GST pull-down assay (Fig. 3A), revealing differential binding compared to RAC1^WT^: weaker binding for RAC1^P29S^, modestly stronger binding for RAC1^F28L^, and no binding for RAC1^T17N^ (Fig. 3B, 3C; supplementary Fig. S7A), which presents representative blots from the GST pull-down assay showing RAC1-PAK1 interactions, with statistical analyses displayed in Figure 3C. Fluorescence polarization further quantified these interactions, confirming no binding for RAC1^T17N^, a modest increase in affinity for RAC1^F28L^, and a 7.5-fold decrease in binding affinity for RAC1^P29S^ relative to RAC1^WT^ (Fig. 3D, 3E; supplementary Fig. S7B), which displays dissociation constants (K_d_) derived from titrations of RAC1 mutants with GST-PAK1 RBD. The slightly enhanced affinity of RAC1^F28L^ was attributed to its slower dissociation rate. Stopped-flow fluorimetry revealed that RAC1^P29S^ and RAC1^F28L^ exhibited 10- and 40-fold slower association rates, respectively, compared to RAC1^WT^, while RAC1^F28L^ displayed a 66-fold and 34-fold slower dissociation rate compared to RAC1^WT^ and RAC1^P29S^, respectively (Fig. 3F, 3G; supplementary Fig. S7C), which provides kinetic analyses of RAC1-PAK1 interactions, including association and dissociation rate constants. Overall, the binding affinity to PAK1 RBD increased slightly for RAC1^F28L^ and decreased 5-fold for RAC1^P29S^ compared to RAC1^WT^ (Fig. 3G), consistent with the results of GST pull-down and fluorescence polarization assays (Fig. 3C, 3E).

**Figure 3.**
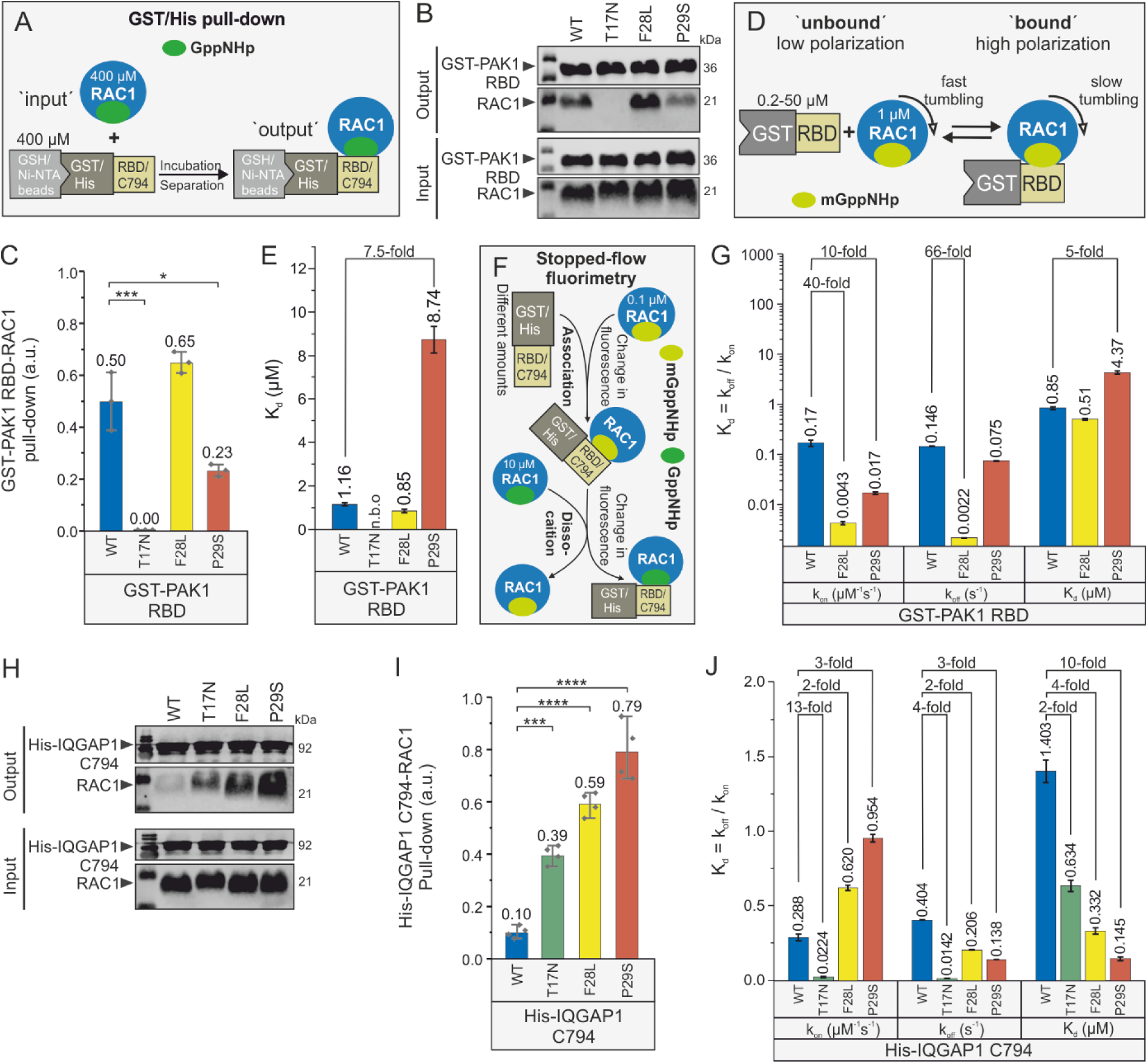
Reduced binding affinity of RAC1^P29S^ for PAK1 but increased for IQGAP1. (**A**) GST and His pull-down assays were performed to evaluate the binding strength of RAC1 variants to GST-PAK1 RBD and His-IQGAP1 C794, respectively. For each reaction, 50 µL of beads were incubated with 400 µM RAC1 proteins and 400 µM GST-PAK1 RBD or His-IQGAP1 C794. Input samples consisted of the protein mixtures before incubation, while output samples were the eluted fractions. (**B**) Western blot analysis of RAC1-PAK1 pull-down (output) was performed using anti-GST antibodies for GST-PAK1 and anti-RAC1 antibodies, with molecular weights indicated in kilodaltons (kDa). The input represents total protein mixtures before pull-down experiments. (**C**) Bar graphs quantify RAC1-PAK1 RBD interactions from 3 independent pull-down experiments analyzed using one-way ANOVA, with P values (* < 0.05; ** < 0.01; *** < 0.001; **** < 0.0001) and data expressed as means ± SD. (**D**) The principle behind the fluorescence polarization measurements for the interaction between GST-PAK1 RBD and RAC1 proteins is illustrated. Accordingly, 1 µM mGppNHp-bound RAC1 was titrated with increasing concentrations of GST-PAK1 RBD. (**E**) Bar graphs from fluorescence polarization analysis represent the dissociation constants (K_d_) for PAK1 RBD binding to RAC1 proteins, with “n.b.o” indicating no binding observed and data expressed as means ± SD. (**F**) The principle behind the kinetic measurements of GST-PAK1 RBD association with and dissociation from RAC1 proteins is shown using a stopped-flow instrument. In these experiments, 0.1 µM mGppNHp-bound RAC1 was rapidly mixed with increasing concentrations of GST-PAK1 RBD to monitor association kinetics. Dissociation kinetics were measured by rapidly mixing a complex of RAC1•mGppNHp•GST-PAK1 RBD with excess GppNHp-bound RAC1. (**G**) Bar graphs from the stopped-flow analysis display the evaluated association rates (k_on_), dissociation rates (k_off_), and dissociation constants (K_d_, calculated as k_off_/k_on_) for the PAK1 RBD interaction with RAC1 proteins, with data presented as means ± SD. (**H**) Western blot analysis of RAC1-IQGAP1 pull-down (output) was performed using anti-His antibodies for His-IQGAP1 and anti-RAC1 antibodies, with molecular weights indicated in kilodaltons (kDa). The Input represents total protein mixtures before pull-down experiments. (**I**) Bar graphs quantify RAC1-IQGAP1 C794 interactions from 4 independent pull-down experiments analyzed using one-way ANOVA, with P values (* < 0.05; ** < 0.01; *** < 0.001; **** < 0.0001), and data expressed as mean ± SD. (**J**) Bar graphs from the stopped-flow analysis depict the k_on_ and the k_off_ values for the interaction between IQGAP1 C794 and RAC1 proteins, with K_d_ values calculated as the ratio of k_off_ to k_on_ and all kinetic data presented as means ± SD.

The interaction of IQGAP1 C794 with RAC1 variants was assessed using a His-tag pull-down assay (Fig. 3A). Binding progressively increased in the order of RAC1^WT^, RAC1^T17N^, RAC1^F28L^, and RAC1^P29S^ (Fig. 3H; supplementary Fig. S8A), which presents representative blots from the pull-down assay showing RAC1-IQGAP1 interactions, with statistical analyses displayed in Figure 3I. This trend was confirmed by data from four independent pull-down experiments (Fig. 3I). Stopped-flow experiments further corroborated these findings, revealing a gradual increase in IQGAP1 binding affinity across the RAC1 variants in the same order (Fig. 3J; Supplementary Fig. S8B), which provides kinetic analyses of RAC1-IQGAP1 interactions, including association and dissociation rate constants. Notably, RAC1^P29S^ exhibited significantly higher affinity for IQGAP1 C794 compared to PAK1 RBD, and IQGAP1 C794 bound more tightly to RAC1^T17N^ than to RAC1^WT^, providing new insights into the differential binding properties of RAC1 effectors.

### RAC1^P29S^ is found in its GTP-bound state in serum-starved cells

To evaluate active RAC1 levels under serum stimulation and starvation, RAC1^WT^ and mutants were overexpressed in HEK-293T cells and pulled down in their GTP-bound states using GST-PAK1 RBD, GST-IQGAP1 C794, and GST as a negative control (supplementary Fig. S9), which provides a schematic representation of the pull-down assay used to determine the level of active, GTP-bound RAC1 from HEK-293T cell lysates.

The results showed significantly stronger binding of GTP-bound RAC1 proteins to IQGAP1 C794 compared to PAK1 RBD under serum-stimulated conditions (Fig. 4A, upper panel). RAC1^P29S^ and RAC1^F28L^ displayed stronger binding to GST-PAK1 RBD, while RAC1^T17N^ exhibited minimal binding relative to RAC1^WT^. In contrast, all RAC1 variants showed significantly higher binding to GST-IQGAP1 C794. Under serum starvation, high levels of RAC1^P29S^•GTP were pulled down with GST-PAK1 RBD, corroborating in vitro findings and indicating temporal accumulation of RAC1^P29S^ in its GTP-bound state (Fig. 4A, lower panel). Similarly, much higher levels of RAC1^P29S^•GTP and RAC1^T17N^•GTP were pulled down with GST-IQGAP1 C794. These findings were reproduced in triplicate, with no interaction observed for GST alone (supplementary Fig. S10), which presents western blot analyses of active GTPase pull-down assays quantifying GTP-bound RAC1 in HEK-293T cell lysates under both serum-stimulated and serum-starved conditions.

**Figure 4.**
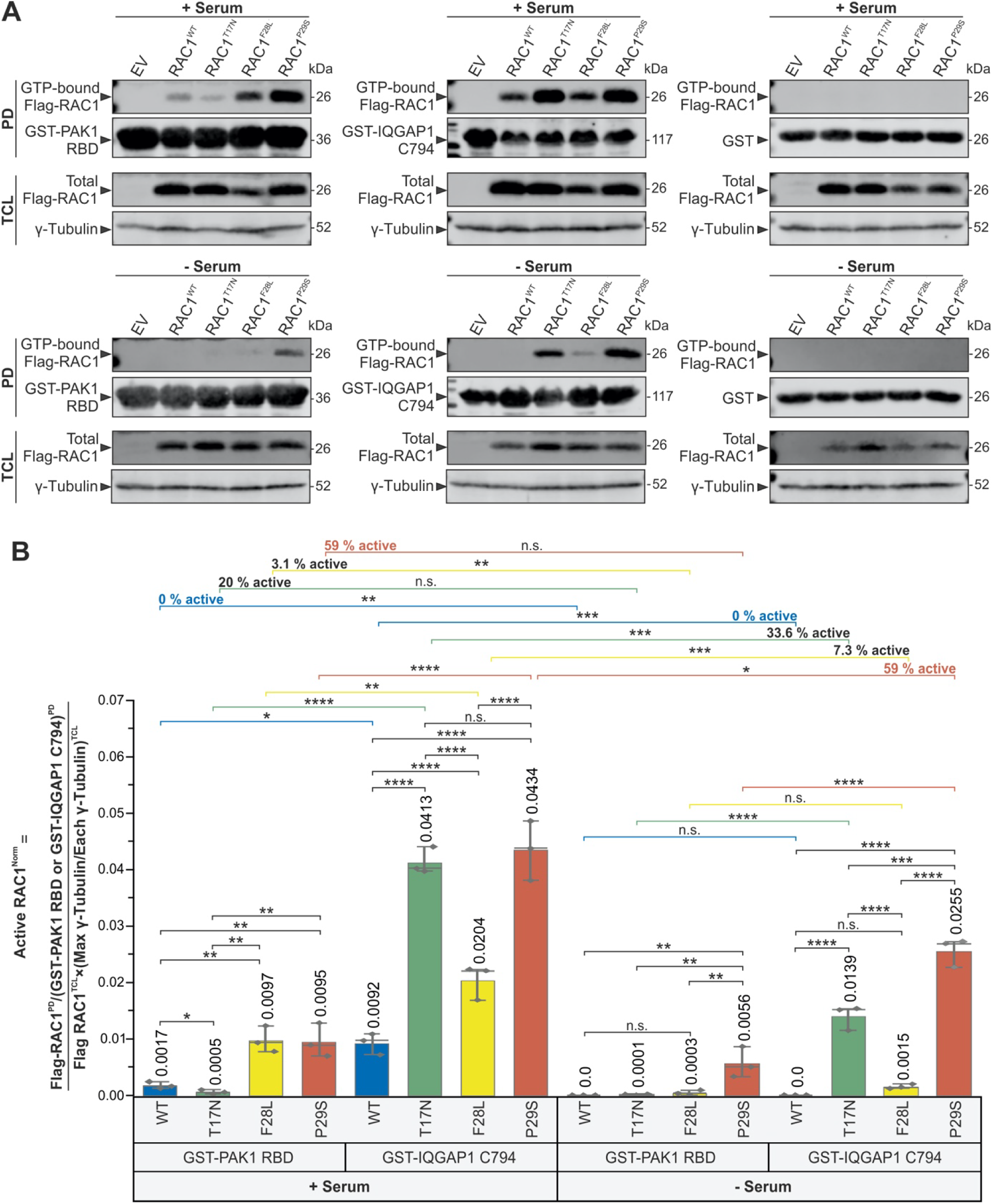
RAC1^P29S^ accumulates in its active, GTP-bound state in HEK-293T cells under serum-starved conditions. Active GTPase pull-down assays were performed to quantify GTP-bound RAC1 proteins (Supplementary Fig. S9). Lysis solutions from *E. coli* containing GST-PAK1 RBD or GST-IQGAP1 C794 were incubated with prewashed glutathione agarose beads to prepare bait-bound beads. Simultaneously, HEK-293T cells were transfected with Flag-RAC1 constructs and cultured under either serum-stimulated or serum-starved conditions for 24 hours. After harvesting, the cells were lysed, and the supernatants containing GTP-bound Flag-RAC1 proteins were collected. Equal amounts of HEK cell lysates were incubated with the bait-bound beads to facilitate protein-protein interactions. After three washes to remove unbound proteins, active GTP-loaded Flag-RAC1 proteins bound to GST-PAK1 RBD or GST-IQGAP1 C794 were eluted and analyzed by SDS-PAGE and Western blotting. (**A**) Western blots of active GTPase pull-down assays were probed with anti-Flag, anti-GST, and anti-γ-tubulin antibodies to detect GTP-bound Flag-RAC1, GST-PAK1 RBD or GST-IQGAP1 C794, and γ-tubulin, respectively. Analyses were performed under serum-stimulated and serum-starved conditions using GST-PAK1 RBD, GST-IQGAP1 C794, and GST as negative controls. Molecular weights (in kDa) are indicated for each band corresponding to the target proteins. The pull-down (PD) lanes show the output signal representing the amount of GTP-bound Flag-RAC1 proteins captured by the bait-bound beads. GST-PAK1 RBD or GST-IQGAP1 C794 bands reflect the amount of bait protein available for RAC1 binding. Total cell lysate (TCL) lanes show Flag-RAC1 expression with γ-tubulin as a loading control. The figure consists of six Western blot panels: the first blot shows the levels of active RAC1^WT^, RAC1^T17N^, RAC1^F28L^, and RAC1^P29S^, with EV indicating the empty vector control. The upper panels show the serum-stimulated condition with GST-PAK1 RBD as bait protein (left panel), GST-IQGAP1 C794 (middle panel), and GST (right panel). The lower panels show the amount of active RAC1 after 24 hours of serum starvation with GST-PAK1 RBD, GST-IQGAP1 C794, and GST from left to right. (**B**) Bar graphs of normalized values from three independent experiments (n = 3), analyzed by one-way ANOVA, were used to quantify active RAC1 proteins. P values are indicated as follows: * < 0.05; ** < 0.01; *** < 0.001; **** < 0.0001, and ns = not significant. Data are expressed as mean ± SD, as detailed in Supplementary Figure 8 (A) and (B). Values for RAC1^WT^, RAC1^T17N^, RAC1^F28L^, and RAC1^P29S^ were compared and analyzed in eight different sets: Set 1 [GST-PAK1 RBD (+serum)], Set 2 [(GST-IQGAP1 C794 (+serum)], Set 3 [(GST-PAK1 RBD (+serum)) vs. (GST-IQGAP1 C794 (+serum))], Set 4 [GST-PAK1 RBD (-serum)], Set 5 [GST-IQGAP1 C794 (-serum)], Set 6 [(GST-PAK1 RBD (-serum)) vs (GST-IQGAP1 C794 (-serum))], Set 7 [(GST-PAK1 RBD (+serum) vs (-serum))], and Set 8 [(GST-IQGAP1 C794 (+serum) vs (-serum))], with the last two sets reporting the percentage of active GTP-loaded RAC1 proteins remaining from serum-stimulated to serum-starved conditions.

Quantification of active, GTP-bound RAC1 levels was performed in three independent experiments for each condition (n = 3; supplementary Fig. S10A and S10B), where separate panels show pull-down results for GST-PAK1 RBD and GST-IQGAP1 C794 in both conditions. Results are presented as bar graphs (Fig. 4B). Under serum stimulation, in set 1 analysis, RAC1^F28L^ and RAC1^P29S^ displayed stronger binding to PAK1 RBD compared to RAC1^WT^, which showed baseline interaction, whereas RAC1^T17N^ demonstrated very weak binding, consistent with its K_d_ values. In set 2, RAC1^T17N^ and RAC1^P29S^ exhibited significantly stronger binding to IQGAP1 C794 compared to RAC1^WT^, which showed baseline interaction, with RAC1^F28L^ demonstrating intermediate binding. Notably, RAC1^T17N^ exhibited binding levels to IQGAP1 C794 similar to RAC1^P29S^. In set 3, all RAC1 variants bound more strongly to IQGAP1 C794 than PAK1 RBD, with RAC1^P29S^ and RAC1^T17N^ showing the highest binding levels.

Under serum starvation, set 4 showed that only RAC1^P29S^ remained active and bound to PAK1 RBD, while RAC1^WT^, RAC1^T17N^, and RAC1^F28L^ showed no significant binding. In set 5, RAC1^WT^ completely lost activity, while RAC1^T17N^ and RAC1^P29S^ remained active and strongly interacted with IQGAP1 C794, although RAC1^F28L^ activity was insufficient to achieve significance. In set 6, RAC1^WT^ lost all activity, failing to bind either PAK1 RBD or IQGAP1 C794. RAC1^T17N^ did not bind PAK1 RBD but bound strongly to IQGAP1 C794, while RAC1^F28L^ showed no significant binding to either effector. RAC1^P29S^, however, is bound more strongly to IQGAP1 C794 than to PAK1 RBD.

Sets 7 and 8 compared RAC1 activity between serum-stimulated and serum-starved conditions. RAC1^WT^ lost all activity under serum starvation, failing to bind either effector. RAC1^T17N^ retained tight binding to IQGAP1 C794 under both conditions, though binding strength decreased by 33.6% under starvation. RAC1^F28L^ lost 93–97% of its binding to PAK1 RBD and IQGAP1 C794, reflecting its fast-cycling nature. In contrast, RAC1^P29S^ retained 59% of its activity under serum starvation, binding strongly to PAK1 RBD and IQGAP1 C794, highlighting its constitutive gain-of-function properties.

### RAC1^P29S^ accumulates in its GTP-bound state in cells and hyperactivates cancer-related signaling pathways

To investigate the impact of RAC1 variants on key signaling pathways, HEK-293T cells were transiently transfected with constructs encoding Flag-tagged RAC1 variants. Western blot analysis revealed a significant increase in ERK1/2 phosphorylation (p-ERK1/2) in cells overexpressing RAC1^P29S^ (p < 0.001, ***) and RAC1T17N (p < 0.05, *) (Figure 5). This increase was consistently observed across triplicate experiments (supplementary Fig. S11), which presents western blot analyses of phosphorylation levels of ERK1/2, AKT(S473), AKT(T308), p38 MAPK, and STAT1 α/β in serum-stimulated HEK-293T cells overexpressing RAC1 variants. The observed ERK hyperactivation aligns with its established role in promoting tumor growth and proliferation. RAC1^P29S^ also significantly elevated p38 MAPK phosphorylation (p < 0.01, **) (Figure 5). Phosphorylation of AKT at serine 473 (S473), a target of mTORC2, and STAT1 α/β phosphorylation were statistically significant (p < 0.05, *) but less pronounced compared to ERK and p38 MAPK. AKT phosphorylation at threonine 308 (T308), a PDK1 target, remained non-significant (n.s.).

**Figure 5.**
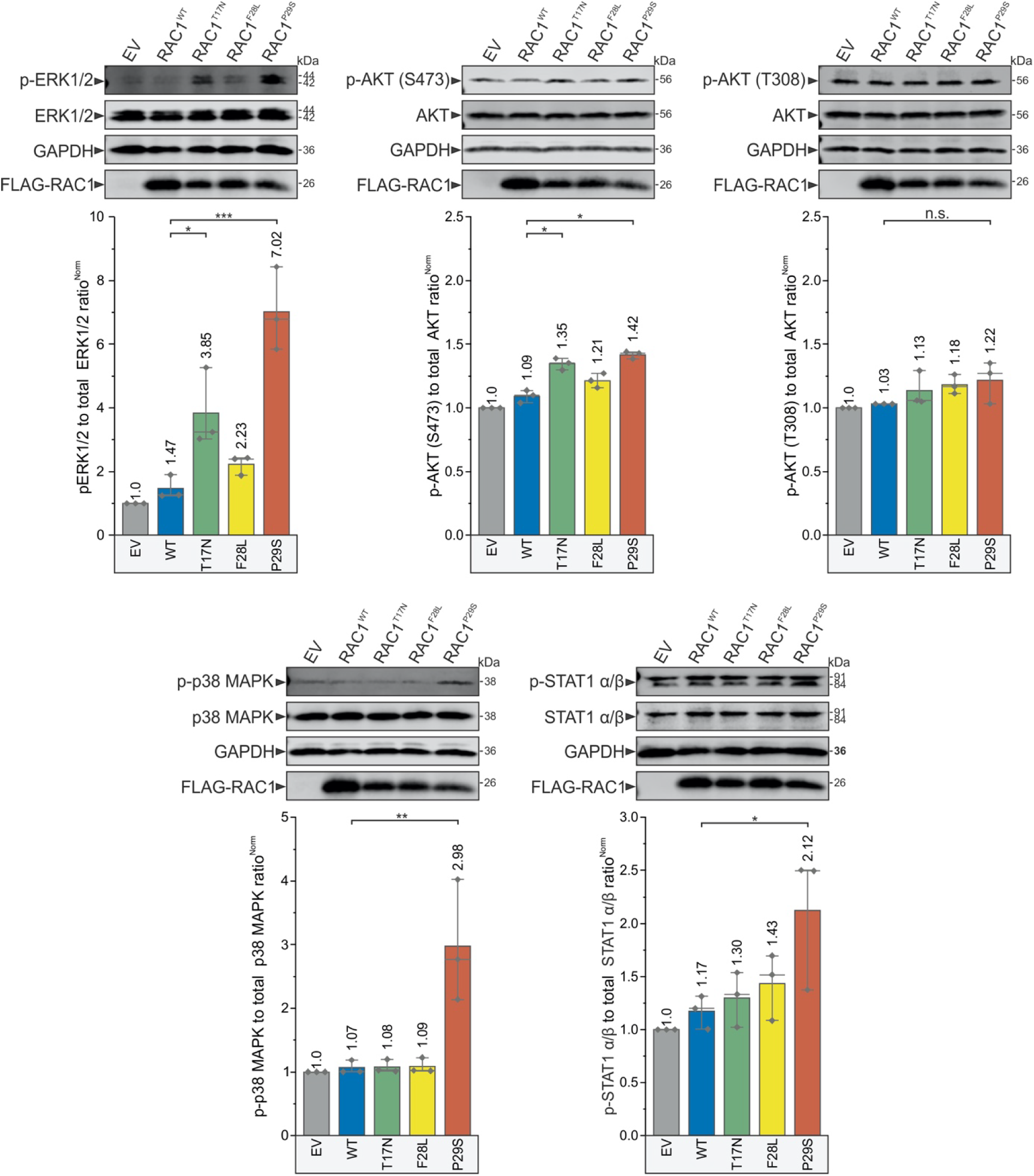
Accumulated GTP-bound RAC1^P29S^ hyperactivates various cancer signaling pathways. Immunoblot analysis was performed to evaluate the phosphorylation levels of several kinases, associated with the hallmarks of oncogenic transformation. Serum-stimulated HEK-293T cells transiently overexpressing Flag-tagged RAC1^WT^, RAC1^T17N^, RAC1^F28L^, and RAC1^P29S^, along with an empty vector (EV) control, were analyzed. The phosphorylation levels of ERK1/2 and AKT (at T308 and S473) were evaluated first. Additionally, the phosphorylation of p38 MAPK was examined as a marker of cellular adaptations that enhance survival under oxidative or inflammatory stress. Finally, the phosphorylation levels of STAT1 α/β, a transcription factor downstream of p38 that may promote immune evasion and support survival under inflammatory conditions, were assessed. Phosphorylation levels were quantified by calculating the ratio of phosphorylated target proteins to total proteins (e.g., p-ERK/t-ERK) and normalizing them to GAPDH as a loading control. Flag tag detection confirmed the expression of each RAC1 variant. Representative results were obtained from three independent experiments (Supplementary Fig. S11), and statistical significance was determined using one-way ANOVA with P values (* P ≤ 0.05; ** P ≤ 0.01; and *** P ≤ 0.001; **** P ≤ 0.0001). Data are expressed as mean ± SD.

## Discussion

This study provides a detailed biochemical characterization of RAC1^P29S^ in comparison to RAC1^WT^, RAC1^T17N^, and RAC1^F28L^ (Fig. 6). Our findings reveal that (i) RAC1^P29S^ exhibits impaired nucleotide binding and accelerated intrinsic nucleotide exchange; (ii) its activation is primarily mediated by DOCK2 rather than DBL family GEFs; (iii) GAP-stimulated GTP hydrolysis is significantly impaired, enabling temporal accumulation of RAC1^P29S^ in its GTP-bound active state; (iv) RAC1^P29S^ exhibits a stronger binding affinity for IQGAP1 compared to PAK1, highlighting IQGAP1 as a spatial modulator of downstream activation; and (v) the accumulation of GTP-bound RAC1^P29S^ leads to hyperactivation of key cancer-associated signaling pathways, including ERK1/2, p38 MAPK. Taken together, these results classify RAC1^P29S^ as a constitutively active mutant and an oncogene (Box 1) that transduces upstream signals to effectors such as IQGAP1, thereby driving processes such as proliferation, invasion, and epithelial-mesenchymal transition (53, 54).

**Figure 6.**
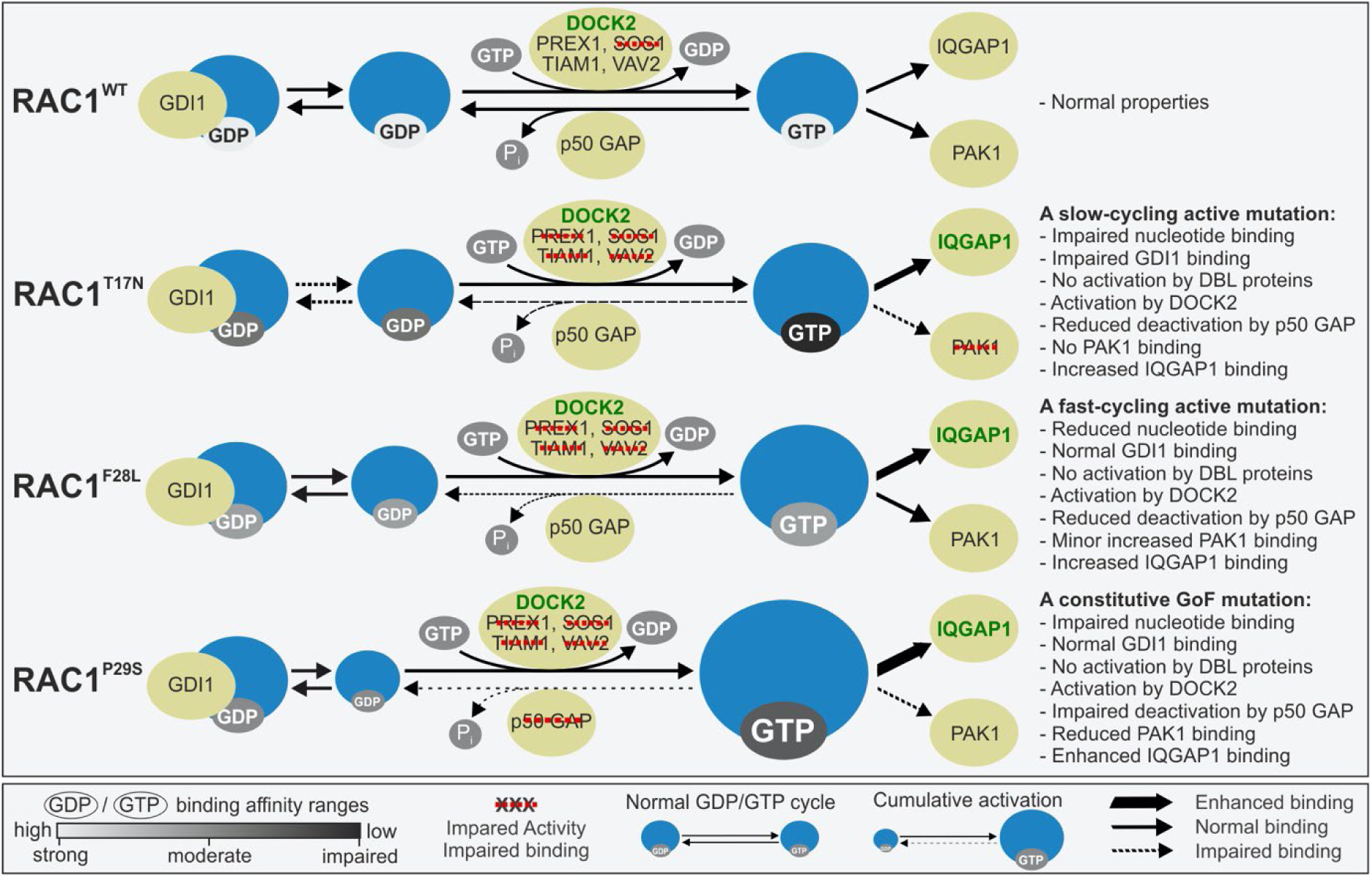
This schematic summarizes the findings of this study, which focuses on the biochemical characterization of RAC1^T17N^, RAC1^F28L^, and RAC1^P29S^ mutants in comparison to RAC1^WT^. The middle section of the figure includes key guides illustrating the strength of GDP/GTP binding, impaired versus enhanced activity or binding to regulators and effectors, and the distinction between the normal GDP/GTP cycle and cumulative activation. Compared to RAC1^WT^, the RAC1^P29S^ mutant significantly impairs nucleotide binding and exhibits a rapid intrinsic nucleotide exchange rate, while the RAC1^T17N^ mutant shows the most impaired nucleotide binding overall. The P29S mutation has a minimal effect on RAC1-GDI1 interaction, whereas the T17N mutation severely impairs GDI1 activity. The P29S mutation is predominantly activated by DOCK2 rather than DBL family GEFs, with GEF-mediated nucleotide exchange being impaired. A key finding of this study is that the P29S mutation significantly impairs GAP-stimulated GTP hydrolysis of RAC1, providing a temporal mechanism for the accumulation of RAC1^P29S^ in its GTP-bound active form and driving its hyperactivation. While the T17N variant shows no binding affinity for PAK1, the P29S mutation demonstrates a dual effect in vitro: reduced binding affinity for PAK1 but enhanced affinity for IQGAP1. This highlights the pivotal role of accessory proteins, particularly IQGAP1, in driving RAC1^P29S^-mediated downstream activation. The rightmost section of the figure provides a detailed summary of the biochemical properties of the RAC1 proteins analyzed in this study.

Our biochemical data confirm that the P29S mutation increases the intrinsic nucleotide exchange rate, consistent with previous reports (36, 37). The slower GDP/GTP association rate results in reduced nucleotide binding affinity despite accelerated exchange. Shimada et al. demonstrated that RAC1P29S enhances GDP dissociation, favoring a GTP-bound state that drives oncogenic activity (55). Similarly, Gursoy et al. used molecular dynamics to show that this mutation increases switch I flexibility, facilitating rapid GDP/GTP exchange (20). Our findings suggest that the elevated exchange rate and activation of RAC1P29S arise from impaired nucleotide-binding affinity due to conformational changes induced by the P29S substitution. However, the intrinsic exchange rate of RAC1^P29S^ remains insufficient for many cellular processes, emphasizing the importance of GEF-mediated exchange in its activation in cancer cells.

RHO-specific GDIs regulate RHO GTPase dynamics by extracting them from membranes, maintaining their inactive state, and preventing degradation through specific interactions (1). Despite progress in understanding GDI-mediated shuttling, some mechanisms remain unclear. We previously showed that GDI1 binds RAC1 regardless of its prenylation state (8). Our data suggest that RAC1^T17N^ has impaired GDI1 activity, with decreased binding affinity, which may suggest persistent plasma membrane association. In contrast, RAC1^P29S^ shows only a slight reduction in GDI1 affinity, indicating that GDI1 can still modulate its localization and translocation.

RAC1^P29S^, like most oncogenes, requires repeated activation by RAC1-specific GEFs. Our data demonstrate that RAC1 mutants exhibit minimal activation by DBL family GEFs, such as TIAM1, PREX1, and VAV2, while DOCK2 significantly enhances the exchange rate for all RAC1 variants, including P29S. This observation aligns with the distinct mechanistic roles of the P-loop and switch I in RAC1, particularly in the functions of DBL and DOCK GEF families (7, 41, 56, 57). However, further analysis is needed to fully understand RAC1^P29S^ activation in cancer cells. Uruno et al. showed that DOCK1 inhibition suppresses cancer cell invasion and macropinocytosis induced by RAC1^P29S^ in melanoma and breast cancer cells (57). Notably, DOCK2 is a potent RAC1 activator in cancers, including melanoma and chronic lymphocytic leukemia (58–60), and regulates critical processes such as lymphocyte migration, T-cell differentiation, cell-cell adhesion, and bone marrow homing of immune cells (61). Although slight increases in TIAM1 activity were observed in our study, the TIAM1-RAC1 axis cannot be entirely excluded from RAC1^P29S^ activation in cancers, including melanoma (62).

RAC1^T17N^ did not show increased GEF-mediated nucleotide exchange via DBL proteins, consistent with its dominant-negative behavior. Overexpression of RAC1^T17N^ in HEK-293T cells significantly increased ERK phosphorylation, though less than RAC1^P29S^. Cool et al. showed that HRAS^D119N^ exhibits dose-dependent dominant-negative and constitutively active effects by reducing nucleotide affinity, sequestering GEFs, binding GTP independently of GEFs, and activating downstream pathways at high concentrations (63). Similarly, RAC1^P29S^ signaling may partially result from overexpression. RAC1^F28L^ shares a GEF activity profile similar to RAC1^P29S^, suggesting both mutations may similarly alter the RAC1 GEF-binding site. This hypothesis requires further structural investigation.

Among the analyzed DBL proteins, SOS1 showed no activity. Other DBL proteins, such as ABR, α-PIX, β-PIX, BCR, FGD4, and FGD6, contain pseudo-DH domains with functions yet to be determined (3). These domains, defined as globular structures performing specific roles like binding or catalysis independent of full-length protein context, may require posttranslational modifications (64–66) or interactions with specific binding partners (67) to become active.

RAC1 signaling is terminated by GTP hydrolysis to GDP, deactivating the protein (5). The intrinsic GTP hydrolysis rate of RAC1^WT^ and its mutants is slow (∼9,000 seconds), necessitating GAPs to catalyze hydrolysis and reduce deactivation time to just a second (48). Our findings confirm that RAC1^P29S^ retains a similar intrinsic hydrolysis rate to RAC1^WT^ (36, 37). However, this study reveals for the first time that the P29S mutation severely impairs GAP-mediated hydrolysis, with p50GAP activity reducing the inactivation time of RAC1^P29S^ to ∼1,000 seconds, a 233-fold decrease compared to RAC1^WT^.

Previous studies have classified RAC1^P29S^ as a spontaneously activating, self-activating, fast-cycling mutant (19, 36, 37) or an oncogenic driver (68) due to its rapid nucleotide exchange that maintains RAC1 in an active state (Box 1). due to its rapid nucleotide exchange that keeps RAC1 active. Our findings align with the latter, highlighting the critical role of p50GAP in regulating RAC1^P29S^ activity. The severe impairment of GAP-stimulated GTP hydrolysis supports its classification as a constitutive gain-of-function mutant and oncogene, driven by defective GAP-mediated deactivation rather than just increased nucleotide exchange. This disruption in temporal regulation leads to the accumulation of active RAC1^P29S^•GTP, as confirmed by its persistence in the GTP-bound state under serum-starved conditions, where most GTPases are typically inactive due to GAP sensitivity and lack of upstream GEF activation. As supported by prior studies, the sustained activation of RAC1^P29S^ likely drives cancer-related processes, including proliferation, survival, invasion, metastasis, and therapy resistance (19, 23–25, 27, 36, 69–71).

The diverse signaling activities of RAC1 are mediated through interactions with specific effectors, which require RAC1 to adopt distinct conformations to function (1). RAC1 effectors include kinases such as PAK1/2/3, MLK1, PI4P5Ks, and accessory proteins like IQGAP1/2, IRSP53, AJUBA, p67phox, and CYFIP1/2 (1). This study examined the binding properties of PAK1, a major kinase, and IQGAP1, a critical scaffolding protein. IQGAP1 is involved in cytoskeletal reorganization processes, including polarity, adhesion, and migration (72, 73), and links RAC1 to the actin cytoskeleton via filamentous actin binding (74). Previous studies showed IQGAP1 interacts with RAC1 and CDC42 via switch regions and effector binding sites, with slight differences in mechanisms (2, 43, 51, 52). Malliri et al. demonstrated that IQGAP1 exhibits increased RAC1 binding specifically upon TIAM1 expression but not other DBL GEFs, including PREX1 (75).

Our findings reveal that RAC1^P29S^ interacts significantly more strongly with IQGAP1 than with PAK1, exhibiting a 30-fold higher binding affinity as measured by stopped-flow fluorimetry. This enhanced interaction was corroborated by a statistically significant increase in RAC1^P29S^•GTP binding to IQGAP1 under both serum-stimulated and serum-starved conditions. In contrast, the stronger binding of RAC1^P29S^ to GST-PAK1 RBD observed in human cell lysates, compared to in vitro pull-down assays using purified proteins, may be attributed to the presence of accessory proteins, modulators, and other cellular components that facilitate protein complex formation in the native environment. These findings suggest that IQGAP1 is a key effector downstream of RAC1^P29S^, acting as an activated scaffolding protein to modulate pathways such as RAF/MEK/ERK (76–78). This underscores the pivotal role of scaffolding proteins, particularly IQGAP1, as spatial modulators facilitating RAC1^P29S^-driven signaling and its downstream effects.

Hyperactivation of signaling pathways downstream of RAC1^P29S^ highlights its oncogenic potential. Accumulated GTP-bound RAC1^P29S^ robustly enhances ERK1/2 and p38 MAPK phosphorylation, suggesting these pathways play significant roles in RAC1^P29S^-driven oncogenic transformation. ERK hyperactivation, a hallmark of uncontrolled tumor growth and proliferation, promotes unregulated cell cycle progression. Concurrently, p38 MAPK hyperactivation supports cellular adaptation to oxidative and inflammatory stress, contributing to tumor progression, invasion, and therapeutic resistance. These findings highlight ERK and p38 MAPK as important mediators of RAC1^P29S^-driven oncogenic signaling, while acknowledging additional pathways may also contribute. Phosphorylation of AKTS473, mediated by mTORC2, and STAT1 α/β, while statistically significant, was less pronounced and may represent secondary or context-specific effects. Selective AKTS473 activation could support cancer cell survival and metabolic adaptation, while STAT1 hyperactivation might facilitate immune evasion and survival under inflammatory conditions. This study focused on these pathways to illustrate GTP-bound RAC1^P29S^ hyperactivation and validate cell-free data highlighting its constitutive activation. However, many other signaling events remain unexplored, underscoring the need for future studies to fully elucidate RAC1^P29S^-driven cancer mechanisms.

## Conclusion

This study highlights the oncogenic potential of RAC1^P29S^ by demonstrating its accumulation in the GTP-bound state and the resulting hyperactivation of downstream signaling pathways (Fig. 6). The P29S mutation significantly impairs nucleotide binding, leading to accelerated intrinsic nucleotide exchange. It is primarily activated by DOCK2 and not by DBL family GEFs. The hyperactivation of RAC1^P29S^ is driven by severely impaired p50GAP-mediated GTP hydrolysis, which serves as a temporal regulatory mechanism facilitating the accumulation of GTP-bound RAC1^P29S^. This accumulation enhances the activation of key cancer-associated signaling pathways, including ERK and p38 MAPK. RAC1^P29S^ exhibits altered binding characteristics that favor IQGAP1 as a critical scaffolding protein in the spatial modulation of downstream signaling. These findings position RAC1^P29S^ as a critical driver of tumorigenesis and suggest that targeting its regulators (DOCK2, p50GAP) and effectors (IQGAP1) may provide promising therapeutic strategies for melanoma.

## Author Contributions Statement

A.M. developed the methods and designed, performed, and analyzed the experiments. A.M. and M.R.A. drafted and approved the final version of the manuscript.

## Acknowledgments

We are grateful to our colleagues at the Institute of Biochemistry and Molecular Biology II for their support, and fruitful discussions.

## Abbreveations

AKT: Protein kinase B
BRAF: v-Raf murine sarcoma viral oncogene homolog B1
DBL: Diffuse B-cell lymphoma
DH-PH: Dbl homology-pleckstrin homology
DHR2: Dock homology region 2
DOCK: Dedicator of cytokinesis
ERK: Extracellular signal-regulated kinase
GAP: GTPase-activating protein
GDP: Guanosine diphosphate
GEF: Guanine nucleotide exchange factor
GDI: Guanine nucleotide dissociation inhibitor
GSH: Glutathione
GST: Glutathione S-transferase
GTP: Guanosine triphosphate
IQGAP1: IQ motif-containing GTPase-activating protein 1
MAPK: Mitogen-activated protein kinase
MLK3: Mixed lineage kinase 3
NRAS: Neuroblastoma RAS viral oncogene homolog
PAK1: p21-activated kinase 1
p50GAP: p50 Rho GTPase-activating protein
PD-L1: Programmed death-ligand 1
PLK1: Polo-like kinase 1
PREX1: Phosphatidylinositol-3,4,5-trisphosphate-dependent Rac exchanger 1
RAC1: Ras-related C3 botulinum toxin substrate 1
RBD: RAC1 binding domain
SOS1: Son of sevenless homolog 1
STAT3: Signal transducer and activator of transcription 3
SUMO: Small ubiquitin-like modifier
TIAM1: T-lymphoma invasion and metastasis-inducing protein 1
VAV2: Vav guanine nucleotide exchange factor 2

## Box 1

The terminologies of the mutations and their effects on the intracellular regulation and function of small GTPases, using the example of RAC1:

- **Dominant negative mutations** impair nucleotide binding affinity to the extent that RAC1 forms a non-functional complex with its cognate GEFs in a nucleotide-free state. A dominant negative RAC1 prevents the activation of wild-type RAC1 when both are present in the same cell, leading to a loss of RAC1 activity and disruption of its downstream signaling pathways.
- **Spontaneous activation mutations** affect the basal activities of RAC1, including enhanced GDP/GTP exchange and reduced GTP hydrolysis. A spontaneously activated RAC1 bypasses normal regulatory mechanisms (without typical regulatory input from other cellular components) and initiates its function independently, leading to unregulated signaling and potentially contributing to cellular dysfunction and disease.
- **Self-activating mutations** refer to the ability of RAC1 to autonomously initiate its signaling function without requiring the usual activation by other cellular components or regulatory proteins. A self-activating RAC1 spontaneously binds GTP and hydrolyzes it to GDP without external regulatory input. This autonomous activation can lead to uncontrolled signaling pathways, potentially contributing to cellular dysfunctions and diseases such as cancer.
- **Fast cycling mutations** lead to the rapid turnover rates of the GDP/GTP exchange and GTP hydrolysis of RAC1. A fast-cycling GTPase rapidly cycles between the inactive, GDP-bound state and the active, GTP-bound state, allowing for quick and dynamic regulation of cellular processes.
- **Constitutively active mutations** affect the GTP hydrolysis reaction of RAC1, resulting in an increased proportion of its active, GTP-bound state, regardless of cellular signaling cues. A constitutively active RAC1 continuously promotes downstream signaling pathways, even when the cell is quiescent or upstream signaling is blocked.
- **Oncogenic mutations** lead to overactivation/hyperactivation of RAC1 and drive oncogenesis. The accumulation of oncogenic RAC1 in its GTP-bound state leads to uncontrolled cellular activities that may contribute to the initiation and progression of various types of cancer.

## Box 2

The definition of various kinetic and equilibrium constants in the context of protein-ligand or protein-protein interactions are described as fallow:

- **Observed rate constants**: The observed rate constant (**k_obs_**) reflects the overall rate at which the interaction occurs, taking into account both the association of the proteins or the protein and ligand, as well as any potential subsequent reactions such as conformational changes or product formation. This is often used in cases where the binding is not at equilibrium and may vary with the concentration of the interacting partners.
- **Association rate constant**: The association rate constant (**k_on_**) measures the rate at which a protein and a ligand or two proteins come together to interact with each other and form a complex. It is defined as: k_on_ = [PL] / [P][L], Where [PL] is the concentration of the protein-ligand or protein-protein complex, and [P] and [L] are the concentration of free protein and free ligand. A higher k_on_ indicates a faster rate of complex formation.
- **Dissociation rate constant**: The dissociation rate constant (**k_off_**) quantifies how quickly the protein-ligand or protein-protein complex dissociates back into the free components. This constant It is important in determining the stability of the interaction; a higher k_off_ indicates a less stable complex. It can be measured experimentally by monitoring the concentration of the complex or over time after dilution or removal of the ligand.
- **Catalytic rate constant**: In the context of enzyme-ligand interactions, the catalytic rate constant (**k_cat_**) refers to the maximum rate of product formation for an enzyme when it is saturated with substrate. For protein-protein interactions, this term may not apply unless there is a specific enzymatic function associated with the interaction, such as in signaling complexes. This constant indicates how efficiently the enzyme catalyzes the reaction after the binding event.
- **Dissociation constants**: The dissociation constant (**K_d_**) is critical for understanding the affinity between a protein and its ligand or between two interacting proteins. It is defined as K_d_ = k_off_ / k_on_. A lower K_d_ indicates a higher affinity between the protein and its partner, meaning they bind more tightly. It is often used to assess the strength of the interaction and is expressed in molar concentration units (M).
- **Equilibrium dissociation constants**: The equilibrium dissociation constant (e**K_d_**) measures the affinity between a protein and a ligand or between two proteins in a more complex or biochemical context. It represents the equilibrium state of a reversible binding interaction and is defined as the ratio of the rate constants of dissociation and association. This constant is particularly relevant when considering interactions that occur in environments where factors such as concentration, binding site availability, or the presence of other interacting partners may influence the overall binding dynamics. In a binding reaction where a ligand (L) binds to a protein (P) to form a complex (PL), It is given by the formula eK_d_ = [R][L] / [RL]. A low eK_d_ value indicates a high affinity between the proteins and its ligand, meaning they bind tightly, while a high eK_d_ value indicates low affinity, meaning they bind weakly.

