## Supplementary Information for "New insights into the classification of the RAC1 P29S hotspot mutation in melanoma as an oncogene"

Amin Mirzaiebadizi[1*], Mohammad Reza Ahmadian[1*]
Institute of Biochemistry and Molecular Biology II, Medical Faculty and University Hospital Düsseldorf, Heinrich Heine University, 40225 Düsseldorf, Germany

### Detailed Material and Methods

**Constructs.** Different pGEX vectors (pGEX-2-T and pGEX-4T-1), encoding an N-terminal glutathione S-transferase (GST) fusion protein, were used to overexpress human RAC1 wild-type (RAC1$^{WT}$: accession number or acc. no. of P63000) and its mutants, including RAC1$^{T17N}$, RAC1$^{F28L}$, and RAC1$^{P29S}$, in *Escherichia coli.* The same expression system was also used to overexpress various regulators and effectors, including human guanosine dissociation inhibitor 1 (GDI1: acc. no. D13989), the Dbl homology-pleckstrin homology (DH-PH) tandem domains of T-lymphoma invasion and metastasis-inducing protein 1 (TIAM1: acc. no. Q13009; amino acids or aa 1033–1404), human vav guanine nucleotide exchange factor 2 (VAV2: acc. no. P52735; aa 168–543), human son of sevenless homolog 1 (SOS1: acc. no. Q07889; aa 189–551), and human phosphatidylinositol-3,4,5-trisphosphate-dependent Rac exchanger 1 (PREX1: acc. no. Q8TCU6; aa 34–415), as well as the GTPase-activating protein (GAP) domain of human p50GAP (acc. no. P85298; aa 198–439), the C-terminal part of human IQ motif-containing GTPase-activating protein 1 (IQGAP1: acc. no. P46940; C794; aa 863–1657), and the CDC42/RAC binding domains (RBD) of human p21-activated kinase 1 (PAK1: acc. no. Q13153; RBD, aa 57–141). In addition, the pET-23b(+) vector was used to overexpress human IQGAP1 (acc. no. P46940; C794; aa 863–1657) as a hexahistidine-tagged protein, and the pOPINS vector was used to express the Dock homology region 2 (DHR2) domain of human dedicator of cytokinesis 2 (DOCK2: acc. no. Q92608; aa 1211–1624) as an N-terminal His6-small ubiquitin-like modifier (SUMO)-tagged protein (1). For eukaryotic expression, all RAC1 constructs with N-terminal tandem decahistidine triple-flag tags were cloned into the pcDNA-3.1 vector.

**Proteins.** All proteins were purified according to previously described protocols (2-6). Briefly, *Escherichia coli* strains, including pLysS BL21(DE3), CodonPlusRIL, and BL21(Rosetta) were transformed to express the target proteins. For protein extraction, cells were incubated at 4 °C with DNase I (10 µg/mL), lysozyme (2 µg/mL), 1% Triton X-100, and a protease inhibitor cocktail (Roche) followed by sonication using a Branson Sonifier S-450A with a 3- to 19-mm titanium probe. Bacterial lysates were centrifuged at 20,000 g for 30 minutes and filtered to obtain soluble fractions. Tagged proteins were purified from the supernatant either as GST fusion proteins using a glutathione-Sepharose column or as His-tagged proteins using Talon cobalt affinity resin. If necessary, the GST tag was cleaved with thrombin (2 units/mg) overnight at 4 °C, followed by a second round of affinity chromatography to remove residual tags. The proteins were then concentrated and pooled after buffer exchange in a buffer containing 30 mM Tris-HCl or HEPES (adjusted to pH 1 unit above or below the isoelectric points of the proteins), 150-250 mM NaCl, 5-10 mM $MgCl_2$, 3 mM DTT, and 0.1 mM GDP for GTPases. Purified proteins were analyzed by SDS-PAGE and Coomassie staining (Supplementary Fig. S1) and stored at -80 °C.

**Preparation of nucleotide-free and fluorescent nucleotide-bound GTPases.** The preparation of nucleotide-free GTPase was performed in two steps, as described (7, 8). Free and bound GDP was degraded using alkaline phosphatase coupled to agarose beads (0.1-1 U per mg of GTPases) in the presence of a 1.5 molar excess of Gpp($CH_2$)p, a non-hydrolyzable GTP analog that is resistant to alkaline phosphatase but sensitive to phosphodiesterase. After GDP was completely degraded and replaced by Gpp($CH_2$)p, snake venom phosphodiesterase (0.002 U per mg of GTPases) was introduced to cleave the nucleotide into GMP, guanosine (G), and inorganic phosphate (Pi). The progress of the reaction was monitored by HPLC using a Beckman Gold instrument equipped with a C18 reversed-phase column and a buffer consisting of 100 mM potassium phosphate (pH 6.5), 10 mM tetrabutylammonium bromide, and 7.5% acetonitrile. After complete degradation of Gpp($CH_2$)p, the sample was centrifuged to remove the beads, snap frozen in liquid nitrogen, and immediately thawed to inactivate the phosphodiesterase. The nucleotide-free GTPases were then aliquoted and stored at −80 °C for subsequent analyses. Fluorescent GDP-bound (inactive) and GppNHp-bound (active) GTPases were prepared by incubating nucleotide-free proteins with a 1.2-fold molar excess of fluorescently labeled nucleotides (methylanthraniloyl or mant-labeled nucleotides: mdGDP and mGppNHp). mdGDP (methylanthraniloyl-2'-deoxy-guanosine 5′-diphosphate) was used instead of mGDP because of its simple monoexponential fluorescence signal, since mGDP equilibrates between the two 2'- and 3'-methylanthraniloyl isomers, resulting in a biphasic fluorescence signal. mGppNHp (methylanthraniloyl guanosine 5'-[beta,gamma-imido]triphosphate) is a non-hydrolyzable analog of GTP. Samples were passed through prepacked NAP-5 columns to exchange the buffer with one free of unbound nucleotides. In addition, GppNHp-bound GTPases were prepared by performing only the first step using alkaline phosphatase and GppNHp, eliminating the need for phosphodiesterase. The concentrations of the nucleotide-bound GTPases were quantified by HPLC under identical buffer conditions, with 25% acetonitrile for the fluorescently labeled nucleotides. The proteins were then stored at −80 °C. The nucleotides were purchased from Jena Bioscience Co.

**Fluorescence kinetic measurements.** Fluorescence-based kinetic measurements, including both long-term and rapid reactions, were performed using a Horiba Fluoromax-4 fluorimeter and a Hi-Tech Scientific stopped-flow spectrophotometer (Applied Photophysics SX20), respectively, as described (6-9). Excitation wavelengths of 355 nm for mdGDP/mGppNHp and 546 nm for tetramethylrhodamine-labeled GTP (tamra-GTP: tGTP) were used for the stopped-flow analysis. Fluorescence detection was facilitated by a photomultiplier equipped with cut-off filters that allowed the detection of wavelengths above 408 nm for mdGDP/mGppNHp and above 580 nm for tGTP. In addition, slow reactions were monitored using the Horiba fluorimeter in combination with quartz cuvettes (Hellma), with an excitation wavelength of 355 nm and an emission wavelength of 446 nm for mant labeled nucleotides.

**Nucleotide-binding assay.** The nucleotide-binding properties of the RAC1 GTPases were assessed using a stopped-flow fluorimetry approach, as described (10). ucleotide association was measured with 0.1 µM fluorescently labeled nucleotides, such as mdGDP and mGppNHp, and varying concentrations of nucleotide-free RAC1 proteins (0.25–5 µM). Measurements were conducted in a buffer containing 30 mM Tris-HCl, pH 7.5, 10 mM $KH_2PO_4$/$K_2HPO_4$, 5 mM $MgCl_2$, and 3 mM DTT, at 25 °C. The association rate constant (kon) was determined by plotting the observed rate constants (kobs) against the concentration of nucleotide-free RAC1 and analyzing the slope via linear regression using Origin software. The dissociation rate constant (koff) of the fluorescent nucleotide from RAC1 proteins (0.1 µM) was measured by adding a 200-fold excess of the non-fluorescent nucleotide (20 µM). The affinity of RAC1 proteins for GDP and GppNHp was calculated by dividing koff by kon, yielding the equilibrium dissociation constants (Kd), as defined in Box 2 of the main text.

**GEF-catalyzed nucleotide dissociation assay.** The rapid GEF-catalyzed nucleotide exchange reaction was monitored by stopped-flow fluorimetry, as described (8). In this assay, 0.1 µM mGDP-bound RAC1 protein in syringe 1 was rapidly mixed with 200-fold excess of non-fluorescent nucleotide (20 µM) and 10 µM of each GEF, including the DH-PH tandem domains of several DBL family members (TIAM1, SOS1, PREX1, and VAV2) and the DHR2 domain of DOCK2, a member of the DOCK family, separately, in syringe 2. The reactions were performed in a buffer containing 30 mM Tris-HCl, pH 7.5, 10 mM $KH_2PO_4$/$K_2HPO_4$, 5 mM $MgCl_2$, and 3 mM DTT, at 25 °C. The observed rate constants were determined by fitting the data to a single-exponential model using Origin software.

**Intrinsic and GAP-stimulated GTP-hydrolysis assays.** The intrinsic GTP hydrolysis rate of RAC1 proteins was determined by HPLC, a reliable method that has been successfully applied to various GTPases, as noted by Eberth and Ahmadian (8). Briefly, the GTPase reaction was initiated by mixing 100 µM nucleotide-free RAC1 with 90 µM GTP in a GAP buffer containing 30 mM Tris-HCl (pH 7.5), 10 mM $KH_2PO_4$/$K2H_PO_4$, 10 mM $MgCl_2$, and 3 mM DTT at 25 °C. Nucleotide contents were monitored by injecting 30 µL of the reaction mixture onto a reverse phase column, with the total volume adjusted to allow for at least seven injections. The reaction was stopped when no further changes in GTP content were observed. The relative GTP content, determined as the ratio [GTP]/([GTP] + [GDP]), was plotted against time, and the data were fitted to a single exponential function using Origin software to calculate the catalytic rate constant ($k_{cat}$) for GTP hydrolysis. GAP-stimulated GTP hydrolysis rates of different RAC1 proteins were determined by stopped-flow fluorimetry using Tamra-GTP, as described (11). A 1 µM solution of nucleotide-free RAC1 was rapidly mixed with 0.8 µM Tamra-GTP in syringe 1, while 1 µM of the catalytic domains of p50 RhoGAP was added in syringe 2, all in the GAP buffer at 25 °C. After reaction initiation, real-time data were recorded and fitted to an exponential function to calculate $k_{cat}$.

**Protein-protein interaction kinetics.** The interaction of RAC1 with GST-GDI1, GST-PAK1 RBD, and His-IQGAP1 C794 was analyzed by stopped-flow fluorimetry to determine $k_{on}$, $k_{off}$, and $K_d$ values, where $K_d$ was calculated by dividing $k_{off}$ by $k_{on}$, as described (6). For the GDI-binding assay, mdGDP-bound inactive RAC1 (0.1 µM) was incubated with varying concentrations of GST-GDI1 (0.25–2 µM). kon values were determined by plotting kobs versus GST-GDI1 concentrations and applying linear regression. koff values were measured by monitoring the release of GST-GDI1 (0.1 µM) from mdGDP-bound RAC1 in the presence of excess non-fluorescent GDP-bound RAC1 (10 µM) using a single-exponential fit. Similarly, mGppNHp-bound active RAC1 (0.1 µM) was used to study interactions with GST-PAK1 RBD and His-IQGAP1 C794. kon values were derived from kobs plots, and koff values were determined in the presence of excess non-fluorescent GppNHp-bound RAC1 (10 µM).

**Fluorescence polarization.** Fluorescence polarization was used to determine the binding affinity between RAC1 and various effector proteins in their three-dimensional state. Experiments were performed using a Fluoromax 4 fluorimeter in polarization mode according to a predefined protocol (6). Gradually increasing concentrations of different effectors were titrated against mGppNHp-bound RAC1 proteins (1 µM) in a buffer containing 30 mM Tris-HCl (pH 7.5), 50 mM NaCl, 5 mM $MgCl_2$, and 3 mM DTT. The total reaction volume was 200 µL, and assays were performed at a constant temperature of 25 °C. Dissociation constants ($K_d$) were determined by fitting the concentration-dependent binding curves to a quadratic ligand binding equation.

**Cell culture and transfection.** HEK-293T cells were cultured in DMEM supplemented with 10% fetal bovine serum (FBS) and 1% penicillin/streptomycin to support optimal growth and prevent contamination. For transfection, RAC1 constructs containing an N-terminal tandem 10 His-triple flag tag were introduced into the cells using TurboFect™ Transfection Reagent (Thermo Fisher Scientific) according to the manufacturer’s protocol. After transfection, cells were harvested under both serum-stimulated and serum-starved conditions. Cell lysis was performed using FISH buffer (50 mM Tris-HCl, pH 7.5; 100 mM NaCl; 2 mM $MgCl_2$; 20 mM β-glycerophosphate; 1 mM $Na_3VO_4$; 10% glycerol; 1x protease inhibitor cocktail (Roche); and 1% IGEPAL (Thermo Fisher Scientific)). Lysis was performed on ice for 10 minutes, followed by centrifugation at 20,000 g for 5 minutes to remove debris. The resulting clear supernatants were used for subsequent analysis, and protein concentrations were determined using the Bradford assay.

***In vitro* pull-down assays.** Pull-down assays were performed to evaluate the ability and strength of RAC1 proteins to bind to PAK1 RBD and IQGAP1 C794. According to the manufacturer's protocols, these assays utilized GST-fused PAK1 RBD with glutathione agarose beads (Macherey-Nagel) and His-IQGAP1 C794 and His Mag Sepharose Ni beads (Cytiva). Briefly, 50 µL of beads per reaction were washed three times and equilibrated in an ice-cold pull-down buffer consisting 50 mM Tris/HCl (pH 7.5), 150 mM NaCl, 10 mM $MgCl_2$, 3 mM DTT, and 5% glycerol. Equimolar ratios of RAC1 protein to GST-PAK1 RBD and His-

IQGAP1 C794 were prepared in separate experiments, with small aliquots used as input controls. The remaining mixtures were incubated with the respective beads for 30 minutes at 4°C under gentle rotation. After three washes with ice-cold buffer, the glutathione-agarose beads were centrifuged at 700 g, while the His-Mag Sepharose Ni beads were magnetically separated using the MagRac6 device (Cytiva). Proteins were eluted from the His Mag pull-down using the buffer supplemented with 500 mM imidazole. The eluates from the His pull-down and protein-bound agarose beads from the GST pull-down were mixed with Laemmli loading buffer (100 mM Tris-HCl, pH 6.8; 33% glycerol; 300 mM dithiothreitol, 6.7% SDS; and 0.01% bromophenol blue), heated at 95°C to denature the proteins, and analyzed for protein-protein interactions using SDS-PAGE and immunoblotting.

**Active GTPase pull-down assay.** A GTPase pull-down assay was performed to assess GTP-bound active RAC1 levels under serum-stimulated and serum-starved conditions in transiently transfected HEK-293T cells (Supplementary Fig. S9), as described (12). Bacterial lysates containing GST-PAK1 RBD and GST-IQGAP1 C794 were prepared separately in 1 mL aliquots, snap frozen, and stored at -80°C. For bead coupling, based on the results of this pre-test, 25 µL of GST-PAK1 RBD lysate and 50 µL of GST-IQGAP1 C794 lysate were incubated with 50 µL of glutathione agarose beads in a total volume of 500 µL of an ice-cold pull-down buffer consisting 50 mM Tris/HCl (pH 7.5), 150 mM NaCl, 10 mM $MgCl_2$, 3 mM DTT, and 5% glyceroat 4°C. After 30 minutes, the beads were washed three times with the buffer and subsequently used for the assay (Supplementary Fig. S10C). HEK-293T lysates were clarified by centrifugation, normalized to equal protein concentrations, and incubated with GST-PAK1 RBD- and GST-IQGAP1 C794- coupled beads in a buffer containing 50 mM Tris-HCl, pH 7.5; 150 mM NaCl; 10 mM $MgCl_2$; 20 mM β-glycerophosphate; 1 mM $Na_3VO_4$; 3 mM DTT; 10% glycerol; protease inhibitor cocktail at 4°C for 20 minutes. The beads were washed three times, resuspended with SDS-Laemmli loading buffer, and heated at 95°C for 7 minutes. Samples were analyzed by SDS-PAGE and immunoblotting

**Antibodies and Immunoblotting.** Primary antibodies were diluted 1:1000 in TBST containing 0.2% Tween-20 and 30% Intercept® (TBS) blocking buffer (Li-Cor). The following antibodies in this study were as followa: α-RAC1 (Merck Millipore, Cat# 05-389), α-6x-His (Thermo Fisher Scientific, Cat# MA5-33032), α-flag (Sigma Aldrich, Cat# F3165), α-γ-tubulin (Sigma Aldrich, Cat# T5326), α-p-ERK1/2 (Thr202/Tyr204) (Cell Signaling Technology, Cat# 4370), α-t-ERK1/2 (Cell Signaling Technology, Cat# 4696), α-p-AKT(S473) (Cell Signaling Technology, Cat# 4060), α-p-AKT(T308) (Cell Signaling Technology, Cat# 2965), α-t-AKT (Cell Signaling Technology, Cat# 2920), α-p-p38 MAPK (Cell Signaling Technology, Cat# 9211), α-p38 MAPK (Cell Signaling Technology, Cat# 8690), α-p-STAT1 (Cell Signaling Technology, Cat# 8826), α-STAT1 (Cell Signaling Technology, Cat# 9176), and α-GAPDH (Cell Signaling Technology, Cat# 2118), α-GST (Cell Signaling Technology #2624). Secondary antibodies, IRDye® 680RD and 800CW donkey anti-rabbit IgG (Cat# 926-68073, 926-32213) and anti-mouse IgG (Cat# 926-68072, 926-32212), were purchased from Li-Cor. Immunoblots were visualized using the Odyssey® XF Imaging System.

**Statistical Analysis.** Data presented in bar graphs of Figures 1, 2, and 3 for all stopped-flow fluorescence measurements represent the average of 3–6 experiments and are expressed as the mean ± S.D. Intrinsic GTP hydrolysis assays (HPLC) and fluorescence polarization experiments were performed in duplicate, with results also expressed as the mean ± S.D. Immunoblot data were analyzed by quantifying the intensities of specific protein bands using Image Studio Lite version 5.2 software. For in vitro pull-down assays, data normalization was performed by dividing the output RAC1 protein by the output effector protein and then dividing this value by the corresponding input RAC1/effector ratio. Input bands were included to ensure the accuracy of the reaction setup. Representative images were obtained from three (supplementary Fig. S7A) or four (supplementary Fig. S8A) experiments. Data for pull-down assays are expressed as mean ± S.D. For active GTPase pull-down assays, normalization was performed according to the equation shown in Fig. 4B. Flag-RAC1 pull-down (PD) values were normalized to the beads-bound GST effector, while overexpressed Flag-RAC1 levels in lysates were normalized to total cell lysate (TCL) levels using γ-tubulin as a loading control. Active RAC1 levels were calculated by dividing the PD values by the corresponding TCL values. Representative images were obtained from three independent experiments (Supplementary Fig. S10A and S10B). Data are expressed as mean ± S.D. For downstream signaling analyses using lysates from transfected HEK-293T cells, data were normalized by calculating the ratio of phosphorylated target proteins to total proteins (e.g., p-ERK/t-ERK), and further normalized to GAPDH signal as a loading control. To avoid false positives, Flag-RAC1 expression levels were excluded from normalization in these experiments. Data were analyzed by one-way ANOVA followed by Tukey's test, with results considered statistically significant at $P \leq 0.05$ (* $P \leq 0.05$; ** $P \leq 0.01$; and *** $P \leq 0.001$; **** $P \leq 0.0001$), Data are expressed as mean ± S.D.

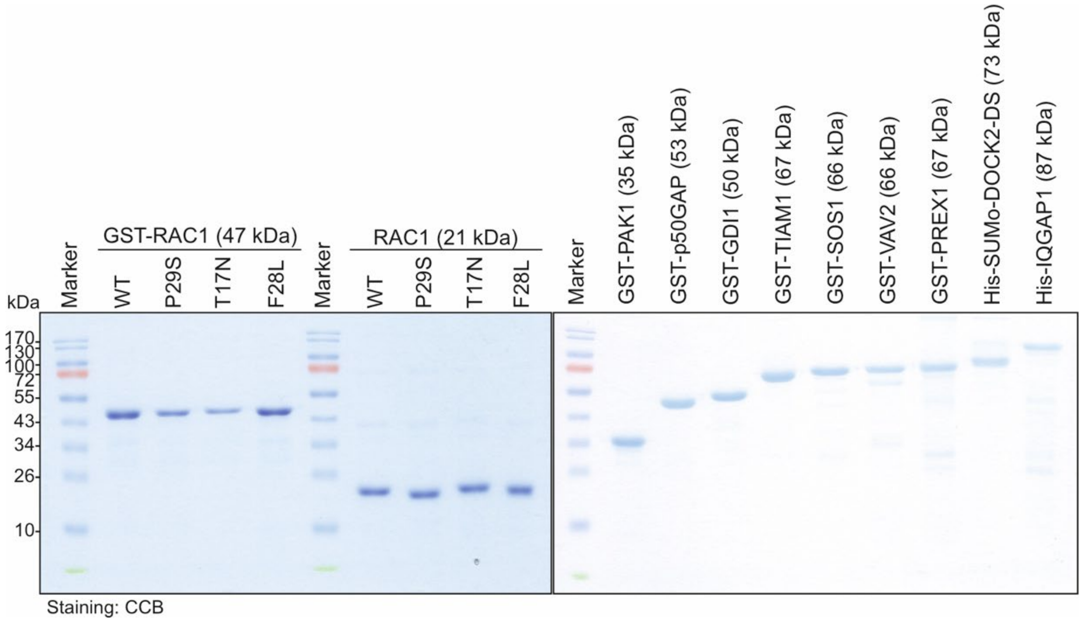


**Figure S1. Purified proteins.** SDS -PAGE gel stained with Coomassie Brilliant Blue (CBB) shows the purified proteins used in this study. For further details, please refer to the Materials and Methods section.

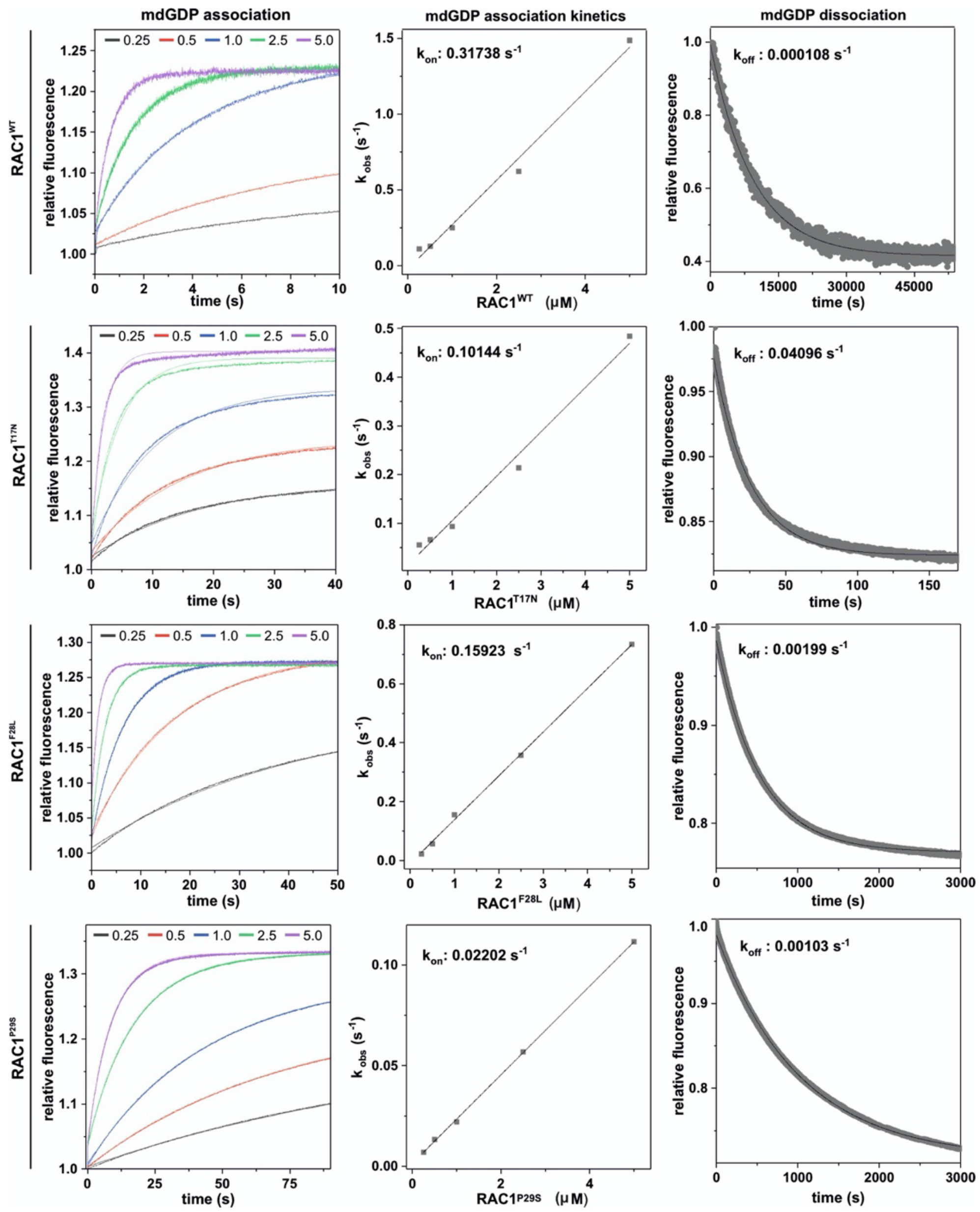


**Figure S2. Kinetic measurements of mdGDP interaction with RAC1 proteins.** The left panels show the interaction of mdGDP (0.1 µM) to RAC1 at increasing concentrations (0.25 to 5 µM). The middle panels illustrate the calculation of the association rate constant ($k_{on}$), derived by plotting the observed rate constants ($k_{obs}$) obtained from the exponential fits of the association data in the left panels against the corresponding RAC1 concentrations, followed by linear fitting. The right panels show the dissociation of mdGDP from RAC1 proteins (0.1 µM) in the presence of excess unlabeled GDP (10 µM). The dissociation rate constants ($k_{off}$) were determined using a single exponential fit. These results are presented as bar graphs in Figure 1.

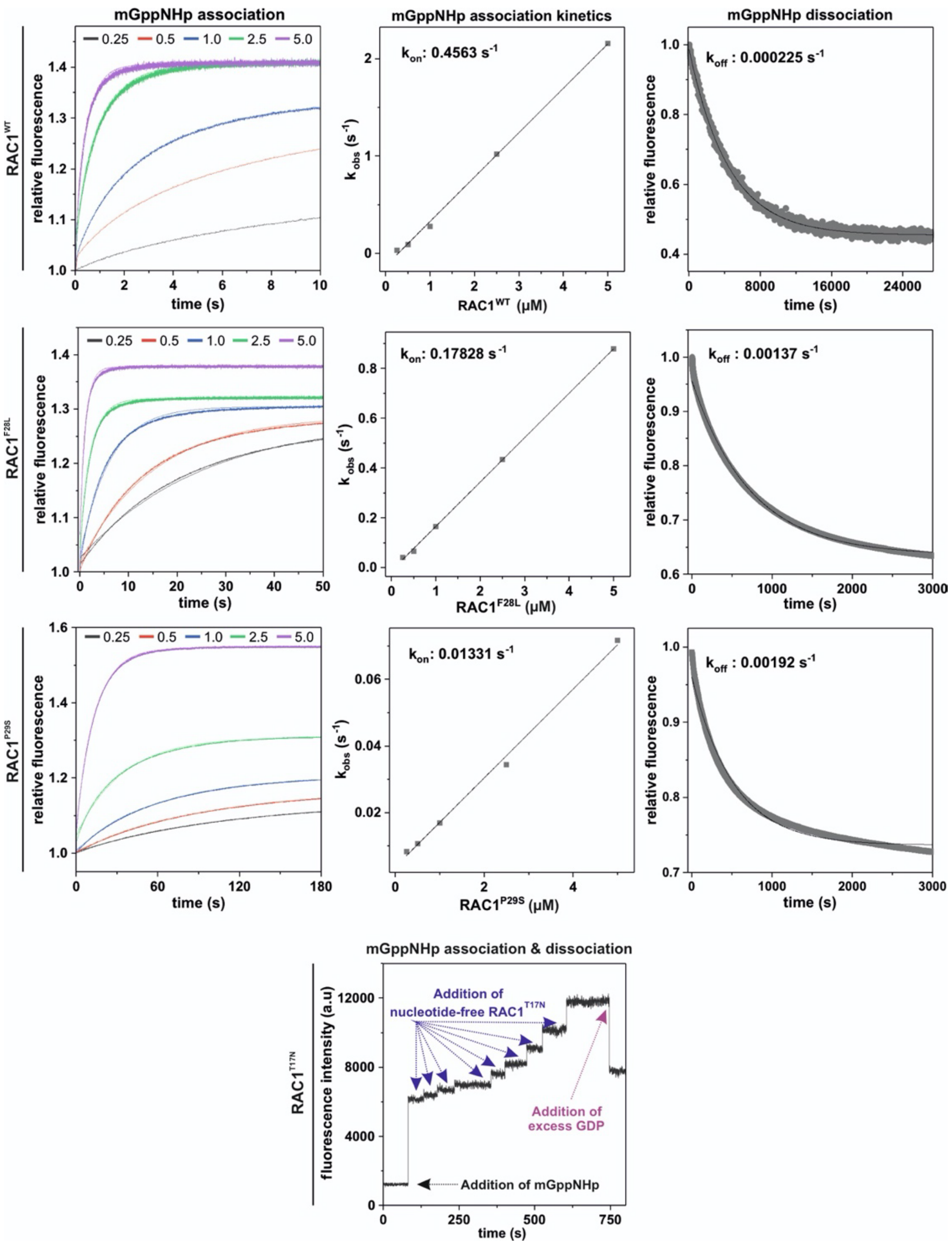


**Figure S3. Kinetic measurements of mGppNHp interaction with RAC1 proteins.** The left panels show the binding of mGppNHp (0.1 µM) to RAC1 over a range of concentrations (0.25-5 µM). In the middle panels, the association rate constant ($k_{on}$) is calculated by plotting the observed rate constants ($k_{obs}$) derived from the exponential fit of the association data in the left panels against the respective concentrations of RAC1, followed by a linear fit. The right panels show the dissociation of mGppNHp from RAC1 proteins (0.1 µM) in the presence of excess unlabeled GppNHp (10 µM). Dissociation rate constants ($k_{off}$) were calculated using a single exponential fit. The data are visually presented as bar graphs in Figure 1. The final lower panel shows fluorescence measurements taken using a standard fluorescence spectrophotometer to investigate the potential association and dissociation kinetics of the mGppNHp interaction with RAC1$^{T17N}$. These measurements could not be performed using stopped-flow fluorimetry. The experiment began with buffer alone, followed by the addition of 1 µM fluorescent mGppNHp, which caused an immediate increase in fluorescence intensity. The association was then assessed by sequentially adding nucleotide-free RAC1$^{T17N}$ in concentrations ranging from 0.05 µM to 1 µM and observing the increases in fluorescence signal. The potential dissociation of mGppNHp from the RAC1$^{T17N}$ was then examined by adding an excess of unlabeled nucleotide (200 µM GDP), which resulted in a rapid drop in signal intensity, indicating dissociation of mGppNHp from nucleotide-free RAC1$^{T17N}$. This experiment demonstrated that the nucleotide-free RAC1$^{T17N}$ can bind to and dissociate from mGppNHp. However, $k_{on}$ and $k_{off}$ values could not be determined using stopped-flow fluorimetry because the association and dissociation rates of nucleotide-free RAC1$^{T17N}$ with mGppNHp were too fast, exceeding the detection capability of stopped-flow fluorescence measurement.

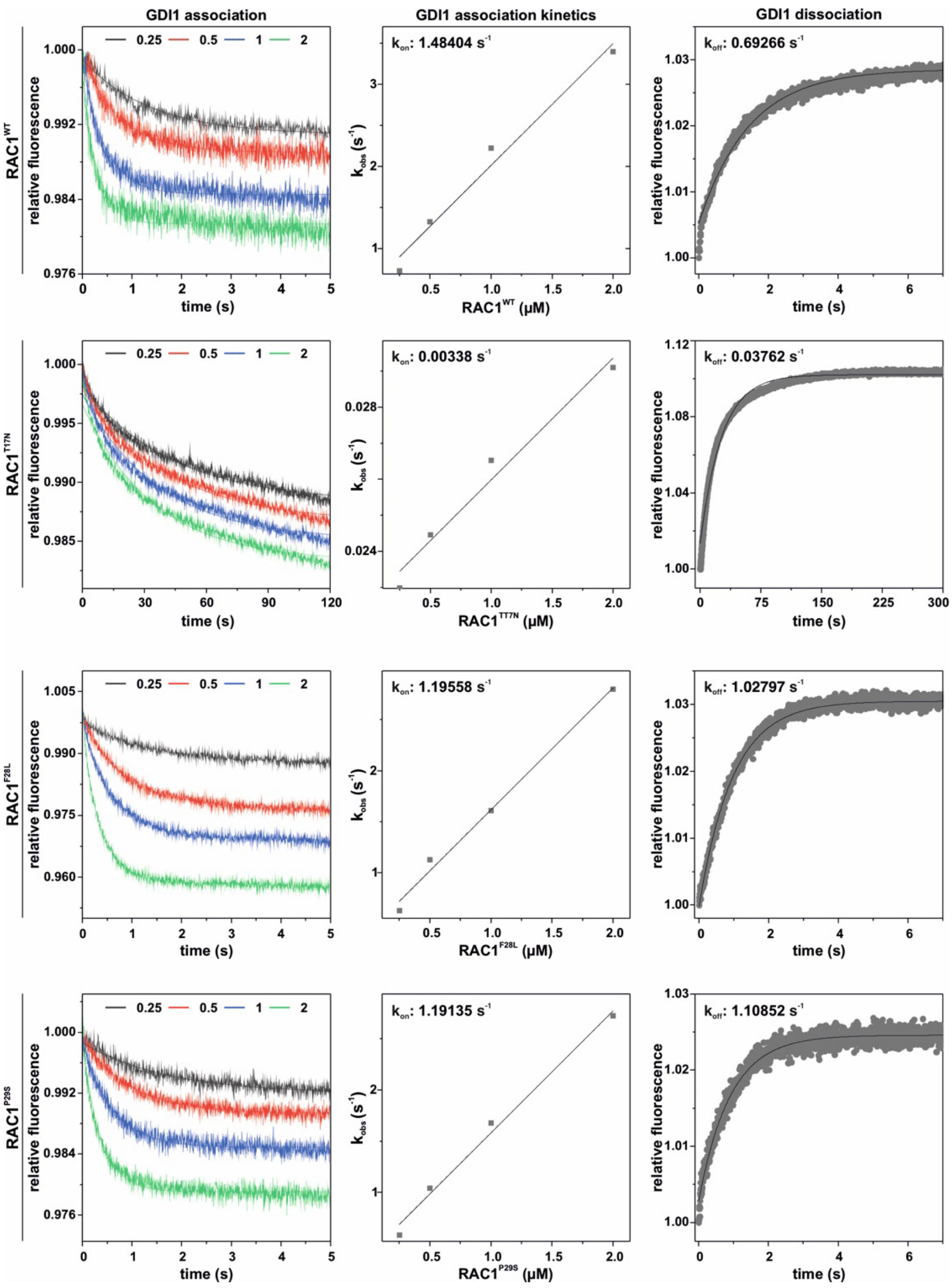


**Figure S4. Kinetic measurements of GDI1 interaction with RAC1 proteins.** The interaction between RAC1 and GST-GDI1 was examined using stopped-flow fluorimetry to determine the association ($k_{on}$) and dissociation ($k_{off}$) rate constants, as well as the dissociation constants ($K_d$) representing the affinity. The left panels show RAC1 binding (0.1 µM) to GST-GDI1 across concentrations ranging from 0.25 to 2 µM. The middle panels illustrate the calculation of $k_{on}$ values, derived by plotting the observed rate constants ($k_{obs}$), obtained from exponential fits of the association data in the left panels, against GST-GDI1 concentrations, followed by linear regression. The right panels depict the dissociation of GST-GDI1 (0.1 µM) from RAC1 proteins (0.1 µM) in the presence of excess non-fluorescent GDP-bound RAC1 (10 µM). $k_{off}$ values were determined through single-exponential fitting. The results are displayed as bar graphs in Figure 2A.

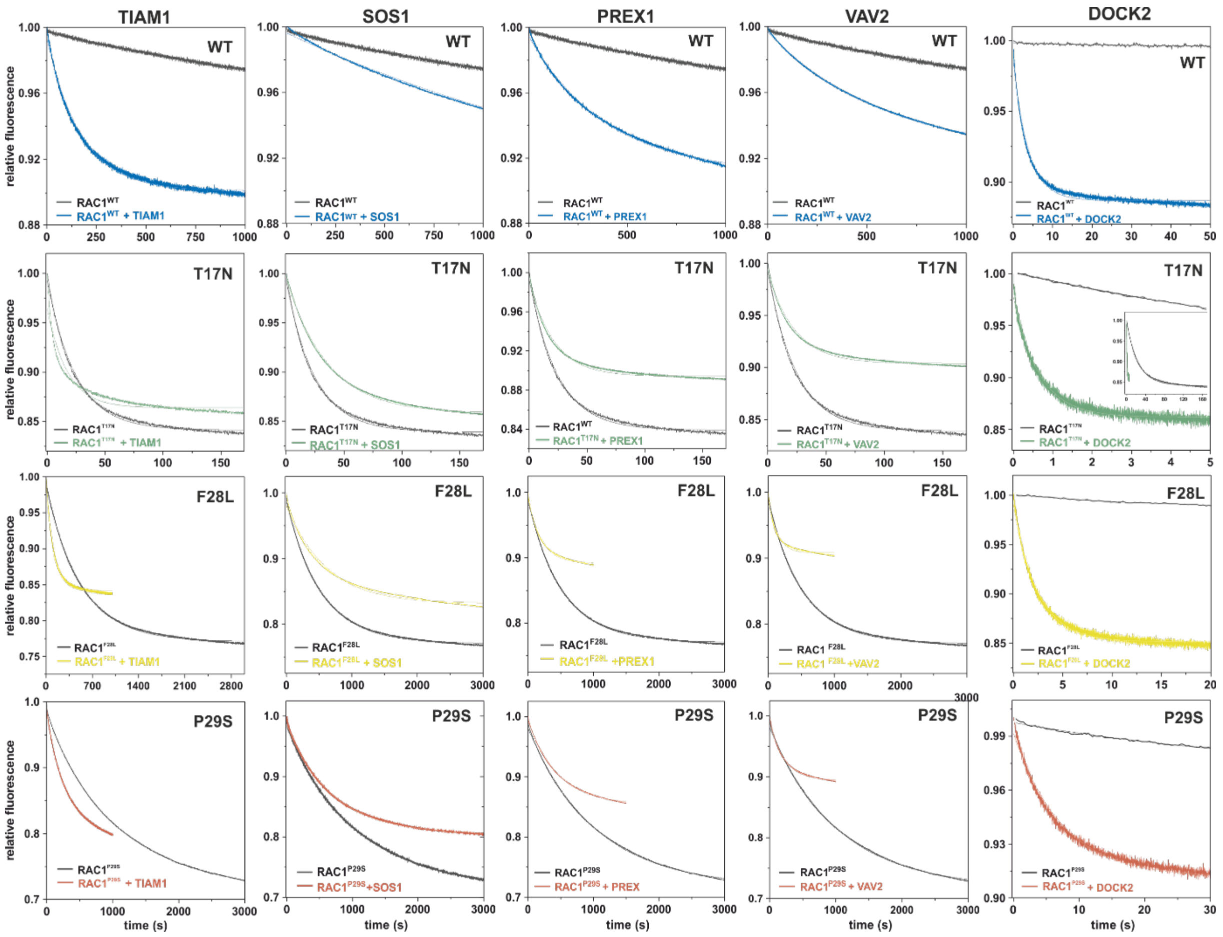


**Figure S5. Kinetic measurements of GEF-catalyzed mdGDP dissociation from RAC1 proteins.** The dissociation of mdGDP from RAC1 proteins (0.1 µM) was measured in the presence of 10 µM GEFs (DH-PH domains of TIAM1, SOS1, PREX1, and VAV2; DHR2 domain of DOCK2) and excess unlabeled GDP (10 µM). The inset show the full kinetics over time. All fluorescence decays were fitted to a single exponential function to calculate the dissociation rate constants ($k_{off}$). The results are presented as bar graphs in Figure 2B.

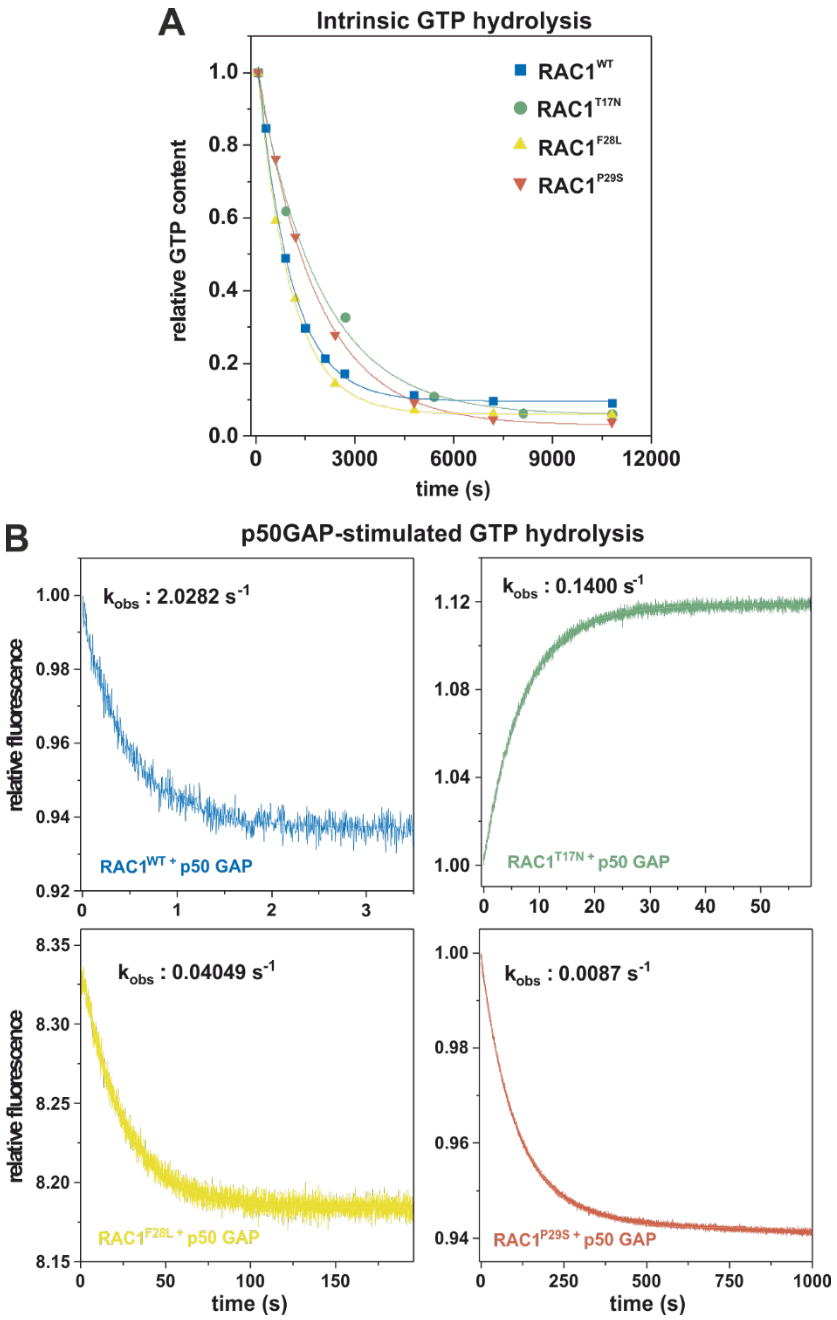


**Figure S6. Measurements of the basal and GAP-stimulated GTP hydrolysis of RAC1 proteins.** (**A**) The basal GTP hydrolysis of the RAC1 proteins was measured using GTP-bound RAC1 proteins and the HPLC method. The relative GTP content, determined as the ratio [GTP]/([GTP] + [GDP]), was used to describe the progress of the reaction. (**B**) Hydrolysis of tGTP (0.8 µM) by RAC1 proteins (0.1 µM) was measured in real-time in the presence of 1 µM p50GAP. All data sets, including HPLC and fluorescence curves, were fitted with a single exponential function to calculate the observed rate constants (kobs), which reflect the catalytic rate constants ($k_{cat}$). The results are presented as bar graphs in Figure 2C.

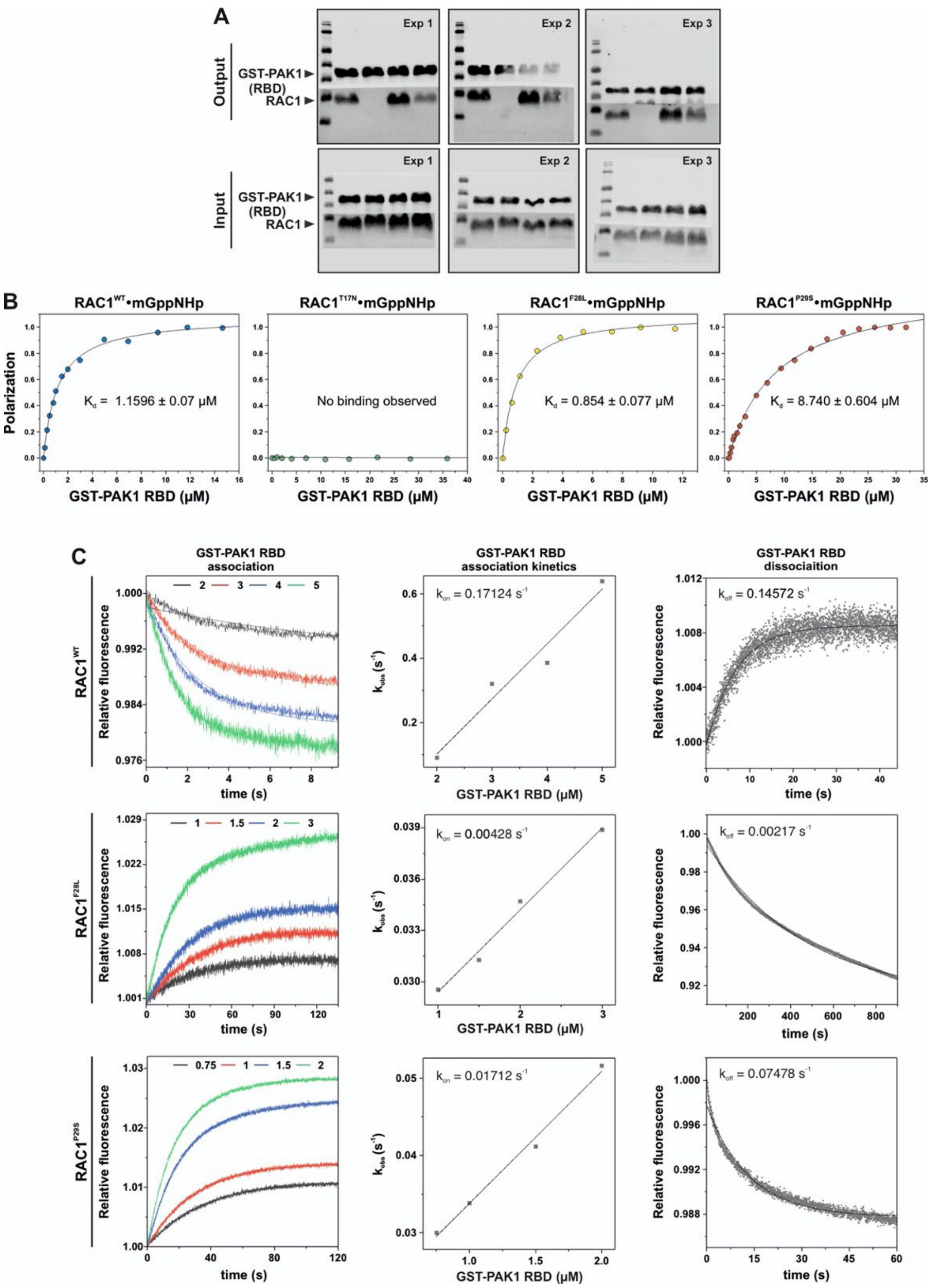


**Figure S7. Measurements of PAK1 interaction with the RAC1 proteins.** (**A**) A GST pull-down assay was performed in triplicate to assess the interaction between RAC1 mutants and GST-PAK1 RBD. The top blots show the pull-down signals (output), while the bottom blots show the input samples before bead incubation. GST-PAK1 RBD was detected with an anti-GST antibody, and RAC1 proteins were visualized with an anti-RAC1 antibody. The input blots confirm that identical amounts of protein were used before bead incubation, ensuring experiment accuracy. A cropped version of experiment 1 is shown in Figure 3B, with data points and statistical analyses displayed as bar graphs in Figure 3C. (**B**) Fluorescence polarization was used to calculate the dissociation constants ($K_d$) for each RAC1 mutant interacting with GST-PAK1 RBD. The mGppNHp-bound RAC1 proteins at 1 μM were titrated with increasing concentrations of GST-PAK1 RBD. Data are shown as bar graphs in Figure 3E. "n.b.o." indicates no binding was observed. (**C**) Stopped-flow fluorimetry was used to assess the affinity of the RAC1 proteins for GST-PAK1 RBD and to provide additional insight into the association ($k_{on}$) and dissociation ($k_{off}$) rate constants, complementing the data from pull-down assays and fluorescence polarization. The left panels show the binding of RAC1 (0.1 µM) to GST-PAK1 RBD at different concentrations (µM). The middle panels show the calculation of the association rate constant ($k_{on}$) by plotting the observed rate constants ($k_{obs}$), derived from the exponential fits of the association data in the left panels, against GST-PAK1 concentrations, followed by linear regression. The right panels illustrate the dissociation of GST-PAK1 RBD (0.1 µM) from RAC1 proteins (0.1 µM) in the presence of excess non-fluorescent GppNHp-bound RAC1 (10 µM). Dissociation rate constants ($k_{off}$) were determined by single-exponential fitting. The results are plotted as bar graphs in Figure 3G.

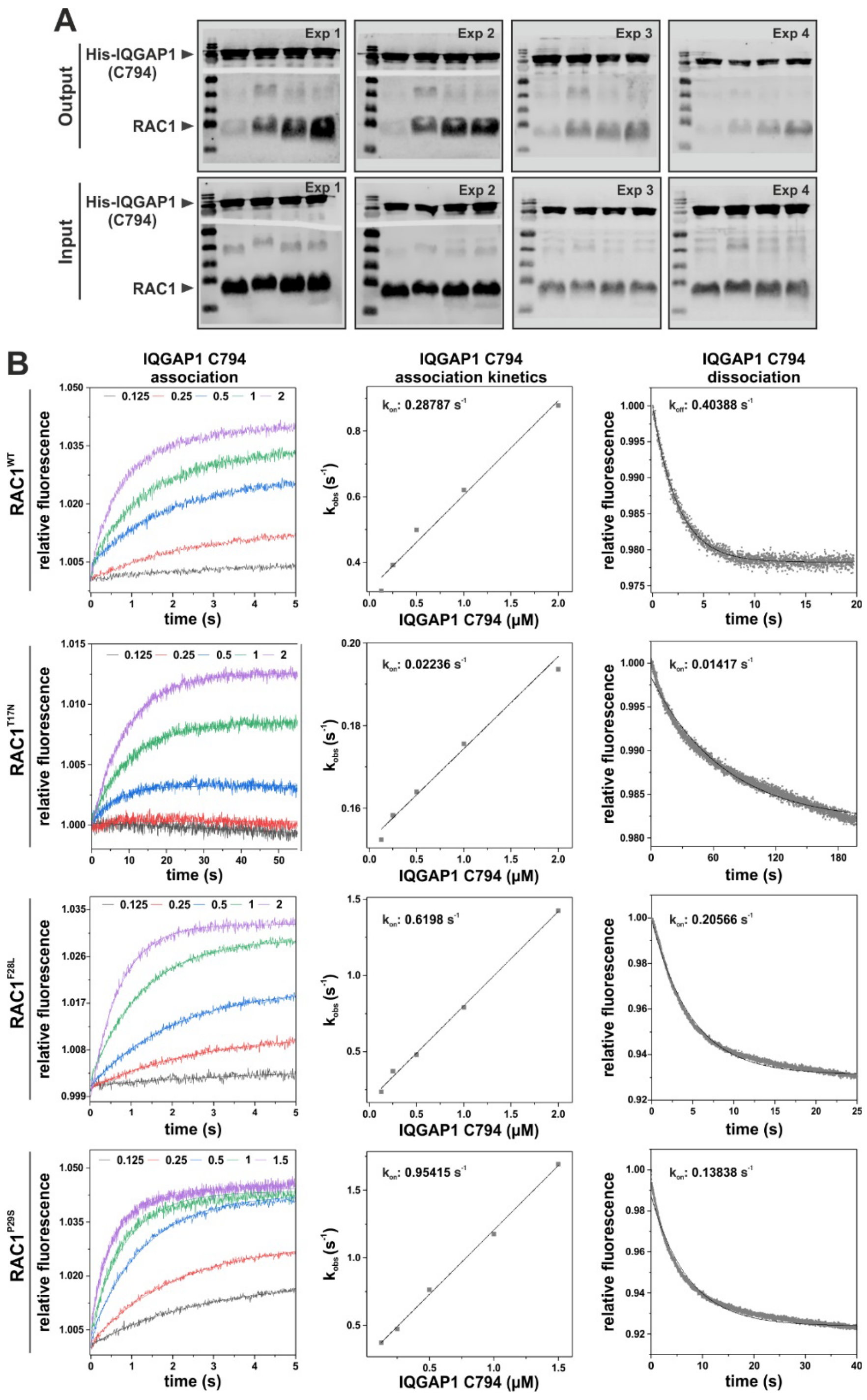


**Figure S8. Measurements of IQGAP1 interaction with the RAC1 proteins.** (**A**) The His pull-down assay was performed in quadruplicate to investigate the interaction between RAC1 mutants and His-IQGAP1 C794. The top blots show the pull-down signals (output), while the bottom blots show the input samples before incubation with His-Mag-Sepharose Ni beads. His-IQGAP1 C794 was detected with an anti-His antibody, and RAC1 proteins were visualized with an anti-RAC1 antibody. The input blots confirm that identical amounts of proteins were used before bead incubation, ensuring the reliability of the results. A cropped version of Experiment 1 is shown in Figure 3H, with data points and statistical analyses presented as bar graphs in Figure 3I. (**B**) Stopped-flow fluorimetry was used to study the interaction between RAC1 and His-IQGAP1 C794 to determine the association ($k_{on}$) and dissociation ($k_{off}$) rate constants, as well as the affinity represented by the dissociation constants ($K_d$). This analysis complements the data obtained from pull-down assays. The left panels show the binding of RAC1 (0.1 μM) to His-IQGAP1 C794 at varying concentrations in μM. The middle panels show the calculation of the $k_{on}$ values, derived by plotting the observed rate constants ($k_{obs}$), obtained from exponential fits of the association data in the left panels, against the concentrations of His-IQGAP1 C794, followed by linear regression. The right panels show the dissociation of His-IQGAP1 C794 (0.1 µM) from RAC1 proteins (0.1 µM) in the presence of excess non-fluorescent GppNHp-bound RAC1 (10 µM). The $k_{off}$ values were determined using single-exponential fitting. The results are plotted as bar graphs in Figure 3J.

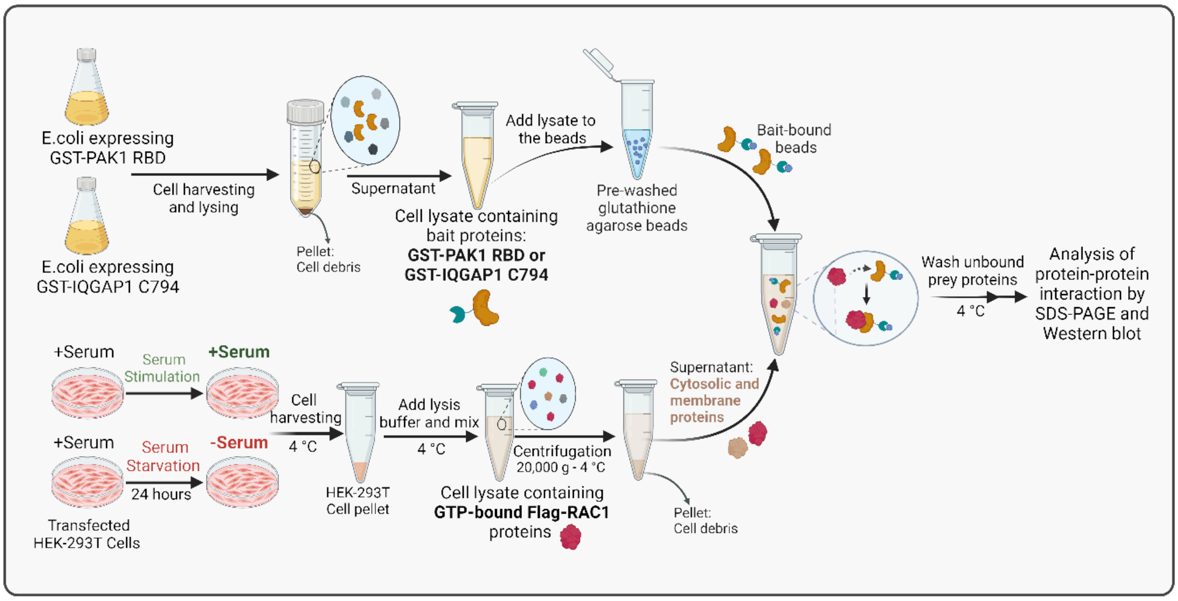


**Figure S9. Schematic representation of the pull-down assay of active, GTP-bound RAC1 from HEK-293T cell lysates.** See the Materials and Methods section for more details. This schematic was created in BioRender.

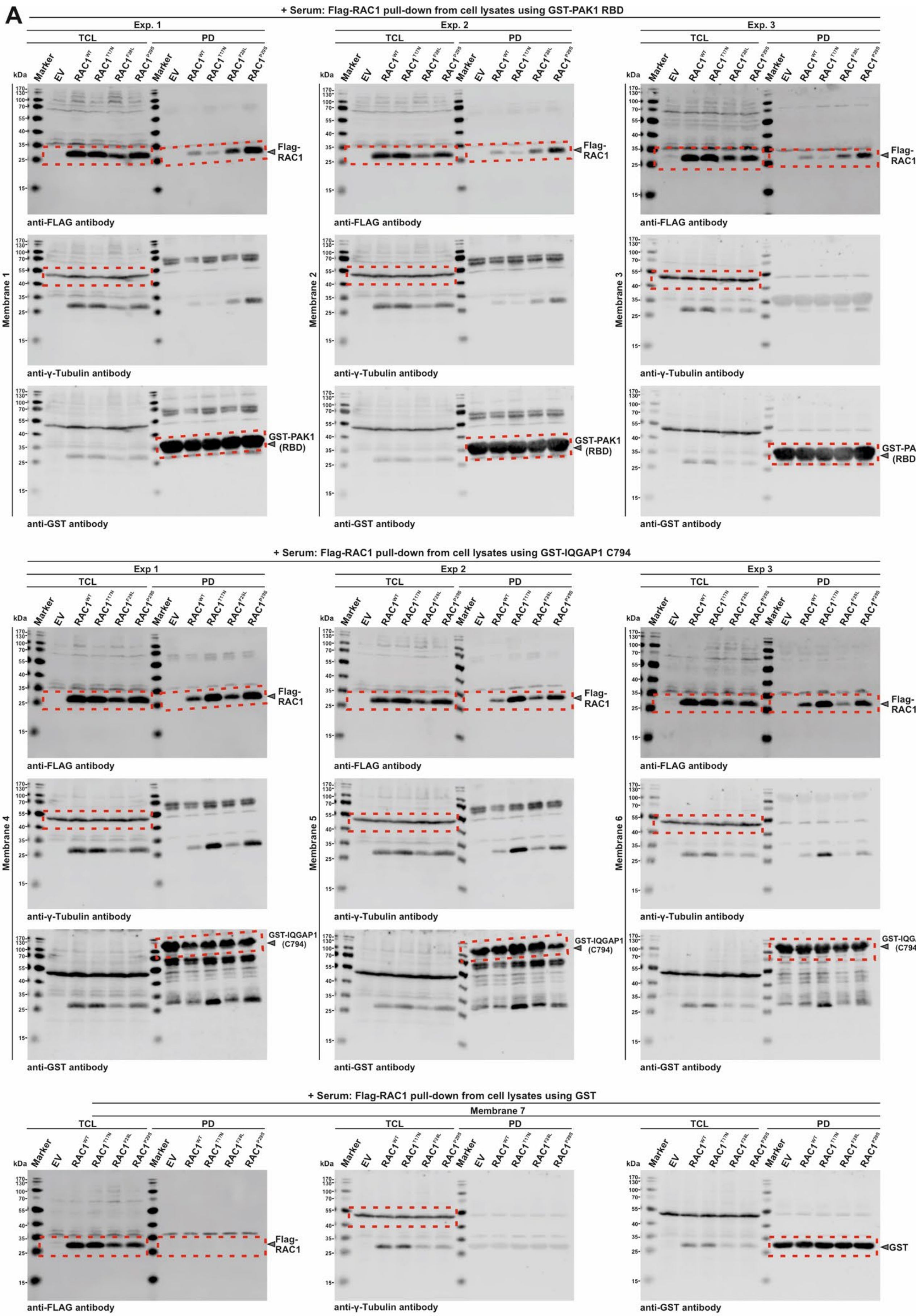
A
+ Serum: Flag-RAC1 pull-down from cell lysates using GST-PAK1 RBD
Exp. 1
Exp. 2
Exp. 3
TCL
PD
Marker
EV
RAC1^WT
RAC1^T17N
RAC1^F28L
RAC1^P29S
kDa
Flag-RAC1
GST-PAK1 (RBD)
anti-FLAG antibody
anti-γ-Tubulin antibody
anti-GST antibody
Membrane 1
Membrane 2
Membrane 3
+ Serum: Flag-RAC1 pull-down from cell lysates using GST-IQGAP1 C794
Exp 1
Exp 2
Exp 3
GST-IQGAP1 (C794)
Membrane 4
Membrane 5
Membrane 6
+ Serum: Flag-RAC1 pull-down from cell lysates using GST
Membrane 7
GST

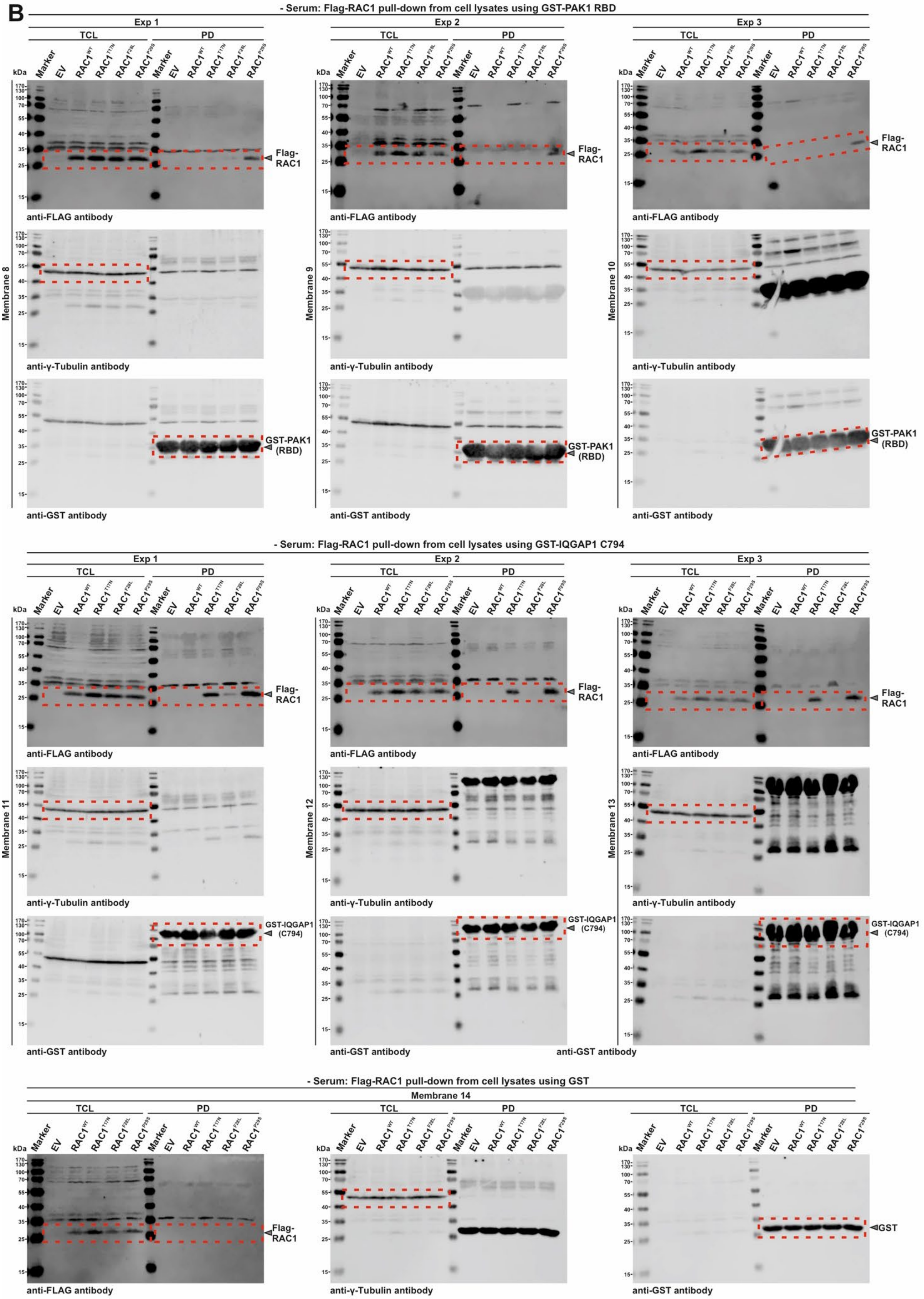
B
- Serum: Flag-RAC1 pull-down from cell lysates using GST-PAK1 RBD
Exp 1
Exp 2
Exp 3
TCL
PD
Marker
EV
RAC1WT
RAC1T17N
RAC1F28L
RAC1P29S
kDa
Flag-RAC1
GST-PAK1 (RBD)
Membrane 8
Membrane 9
Membrane 10
anti-FLAG antibody
anti-γ-Tubulin antibody
anti-GST antibody
- Serum: Flag-RAC1 pull-down from cell lysates using GST-IQGAP1 C794
GST-IQGAP1 (C794)
Membrane 11
Membrane 12
Membrane 13
- Serum: Flag-RAC1 pull-down from cell lysates using GST
Membrane 14
GST

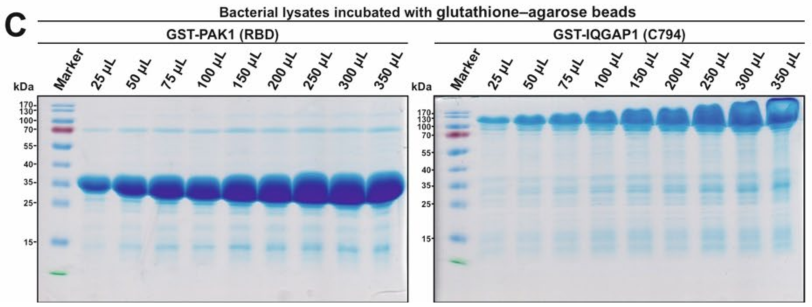


**Figure S10. Active GTPase pull-down assays using GTP-bound Flag-RAC1 from HEK-293T cell lysates.** Active GTPase pull-down assays were performed to quantify GTP-bound Flag-RAC1 from HEK-293T cell lysates with GST-PAK1 RBD, GST-IQGAP1 C794, and GST as negative controls. The assays were conducted under both serum-stimulated (**A**) and serum-starved (**B**) conditions. HEK-293T cells were transfected with RAC1 constructs, serum stimulated or starved for 24 hours, and then harvested and lysed for the pull-down assays. (**A**) shows three panels under serum-stimulated conditions (from top to bottom): the top panel represents GTP-bound Flag-RAC1 pull-down using GST-PAK1 RBD (performed in triplicate), the middle panel represents GST-IQGAP1 C794 (also in triplicate), and the bottom panel shows the negative control, GST, with no interaction observed with Flag-RAC1. (**B**) shows the same three panels under serum-starved conditions. Western blots were probed with anti-Flag, anti-GST, and anti-γ-tubulin antibodies to detect GTP-bound Flag-RAC1, the GST fusion proteins, and γ-tubulin, respectively. Molecular weights (in kDa) are indicated for each protein band. Pull-down (PD) lanes show GTP-bound Flag-RAC1 captured by the bait-bound beads, while total cell lysate (TCL) lanes show Flag-RAC1 expression levels, with γ-tubulin as a loading control. Each membrane contains samples from three independent experiments, labeled Exp1-3, with antibody incubations indicated by red dashed boxes. Membranes 1-3 represent GST-PAK1 RBD under serum stimulation; membranes 4-6, GST-IQGAP1 C794 under serum stimulation; membrane 7, GST under serum stimulation; membranes 8-10, GST-PAK1 RBD under serum starvation; membranes 11-13, GST-IQGAP1 under serum starvation; and membrane 14, GST under serum starvation. (**C**) Coomassie-stained SDS-PAGE Gels showing the bead saturation assay for GST-PAK1 RBD (left) and GST-IQGAP1 C794 (right). After IPTG induction and protein expression, varying amounts of bacterial lysate (25-350 µL) were incubated with 50 µL of GSH beads for one hour and washed three times. The beads were then mixed with SDS-Laemmli sample buffer, heated to 95°C, and analyzed by SDS-PAGE fallowed by Coomassie Brilliant Blue staining. Based on this pre-test, 25 µL of GST-PAK1 RBD lysate and 50 µL of GST-IQGAP1 C794 lysate were used for the active GTPase pull-down assays.

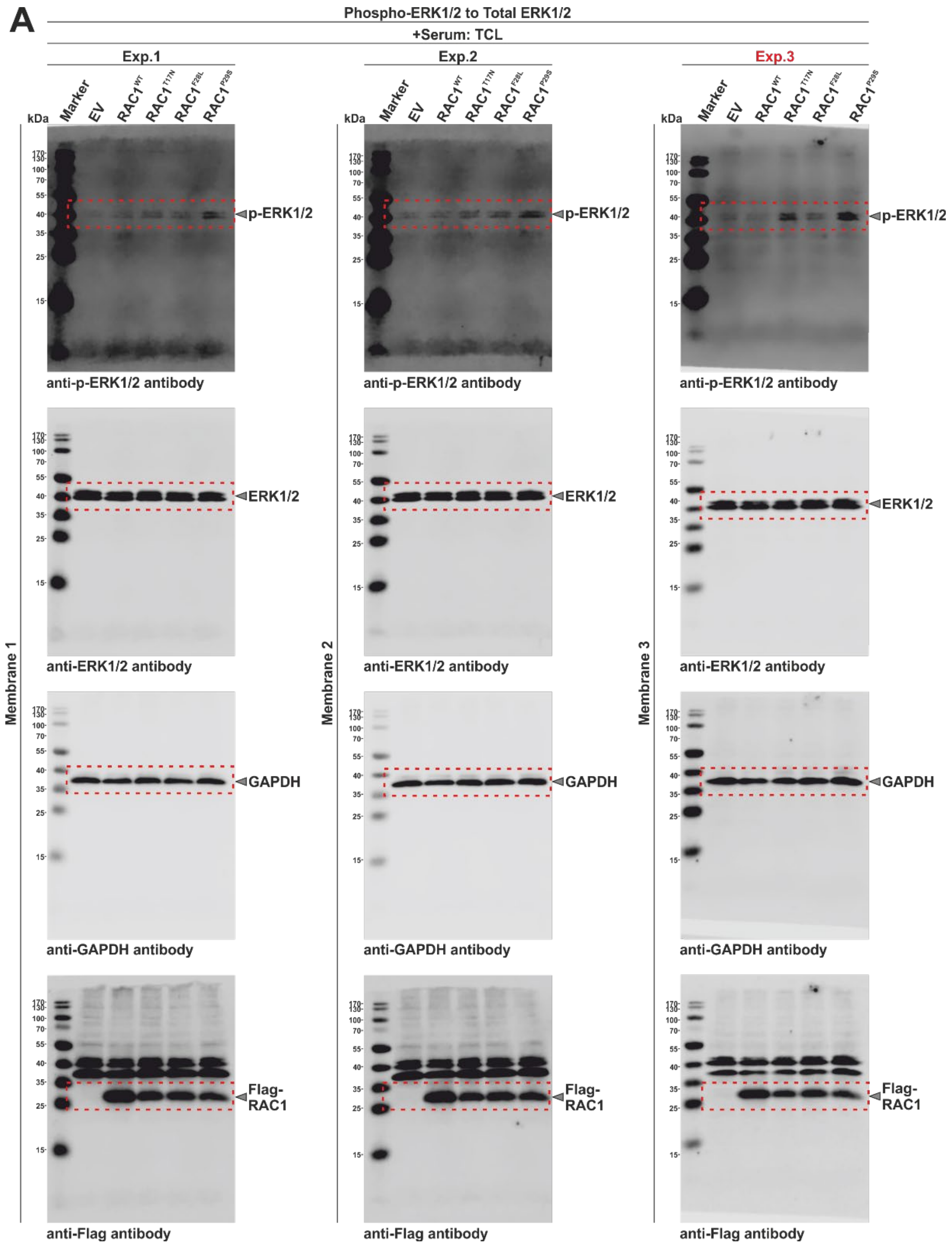

A
Phospho-ERK1/2 to Total ERK1/2
+Serum: TCL
Exp.1
Exp.2
Exp.3
Marker
EV
RAC1WT
RAC1T17N
RAC1F28L
RAC1P29S
kDa
p-ERK1/2
ERK1/2
GAPDH
Flag-RAC1
anti-p-ERK1/2 antibody
anti-ERK1/2 antibody
anti-GAPDH antibody
anti-Flag antibody
Membrane 1
Membrane 2
Membrane 3

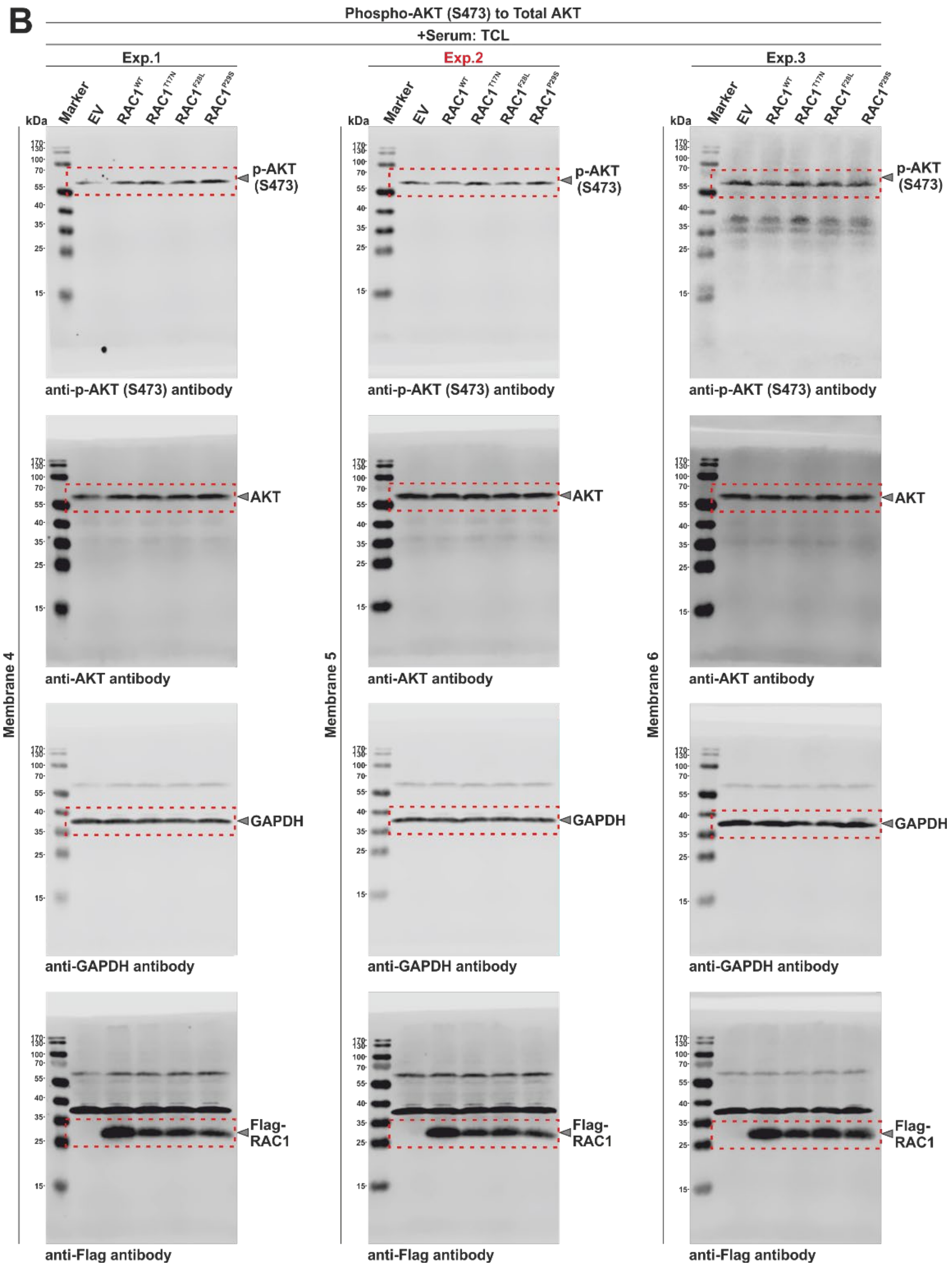
B
Phospho-AKT (S473) to Total AKT
+Serum: TCL
Exp.1
Exp.2
Exp.3
kDa
Marker
EV
RAC1WT
RAC1T17N
RAC1F28L
RAC1P29S
p-AKT (S473)
anti-p-AKT (S473) antibody
AKT
anti-AKT antibody
GAPDH
anti-GAPDH antibody
Flag-RAC1
anti-Flag antibody
Membrane 4
Membrane 5
Membrane 6

C

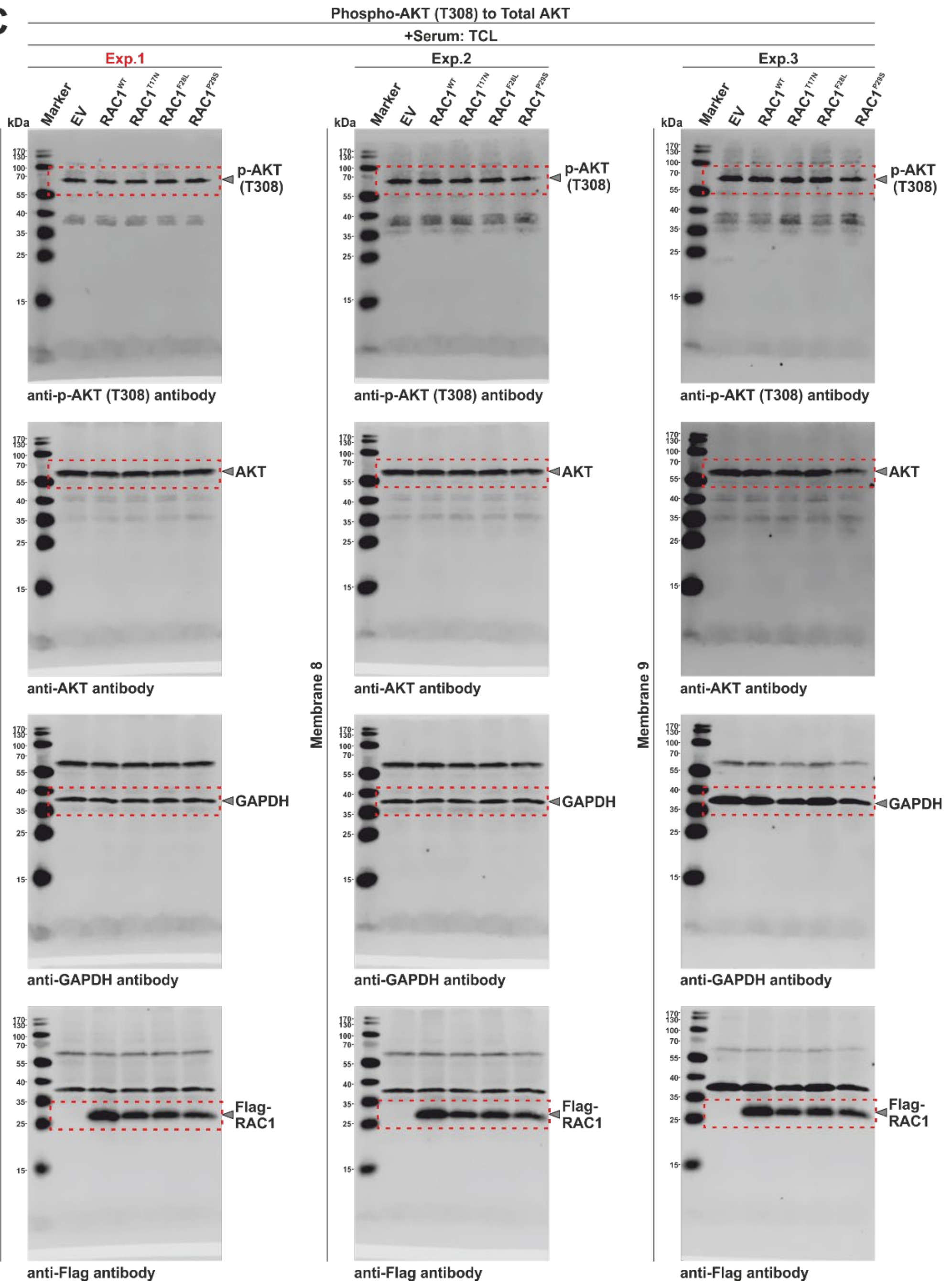
Phospho-AKT (T308) to Total AKT
+Serum: TCL
Exp.1
Exp.2
Exp.3
Marker
EV
RAC1WT
RAC1T17N
RAC1F28L
RAC1P29S
kDa
p-AKT (T308)
AKT
GAPDH
Flag-RAC1
anti-p-AKT (T308) antibody
anti-AKT antibody
anti-GAPDH antibody
anti-Flag antibody
Membrane 7
Membrane 8
Membrane 9

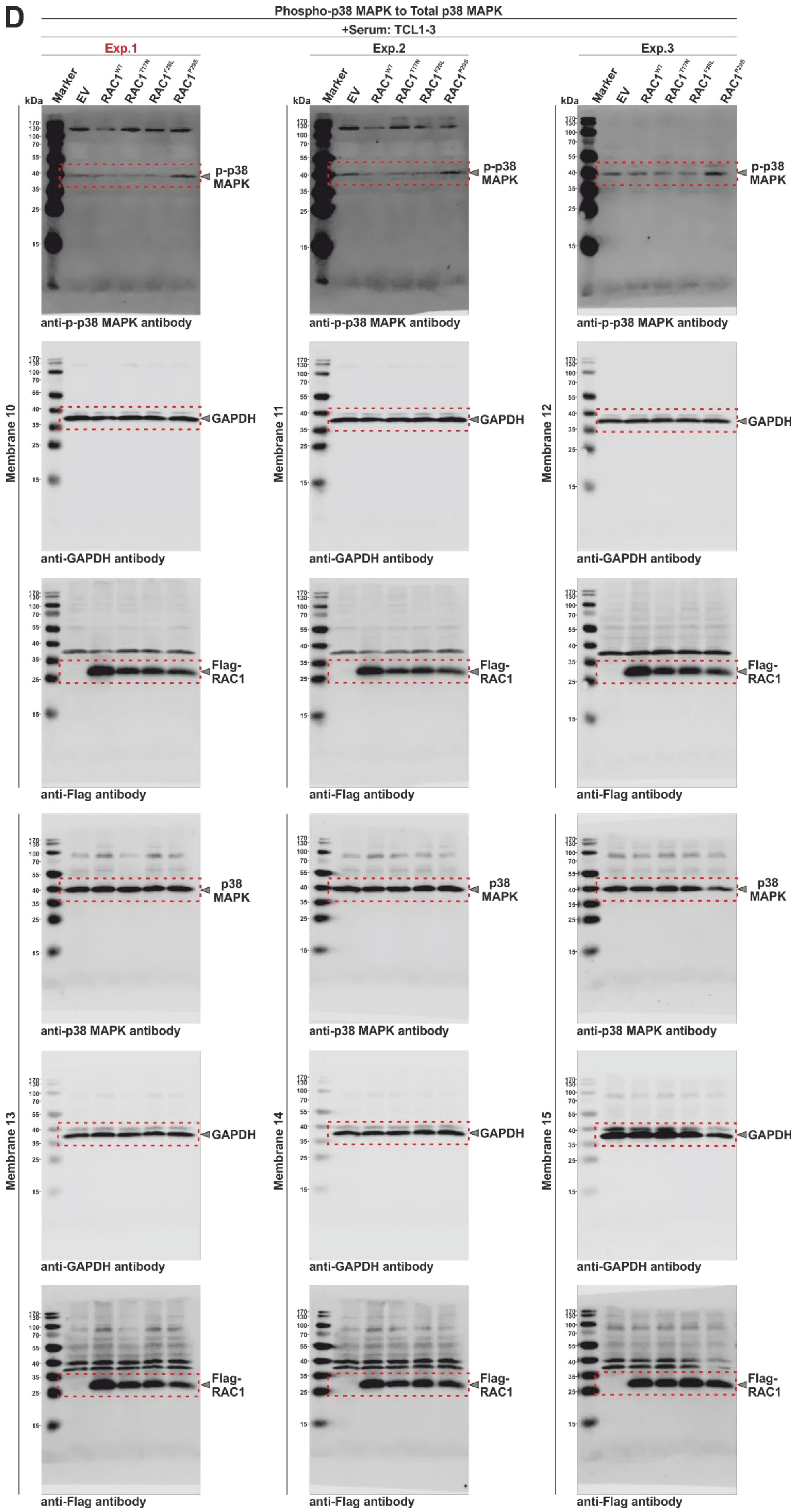
D
Phospho-p38 MAPK to Total p38 MAPK
+Serum: TCL1-3
Exp.1
Exp.2
Exp.3
Marker
EV
RAC1WT
RAC1T17N
RAC1F28L
RAC1P29S
kDa
p-p38 MAPK
anti-p-p38 MAPK antibody
GAPDH
anti-GAPDH antibody
Flag-RAC1
anti-Flag antibody
Membrane 10
Membrane 11
Membrane 12
p38 MAPK
anti-p38 MAPK antibody
Membrane 13
Membrane 14
Membrane 15

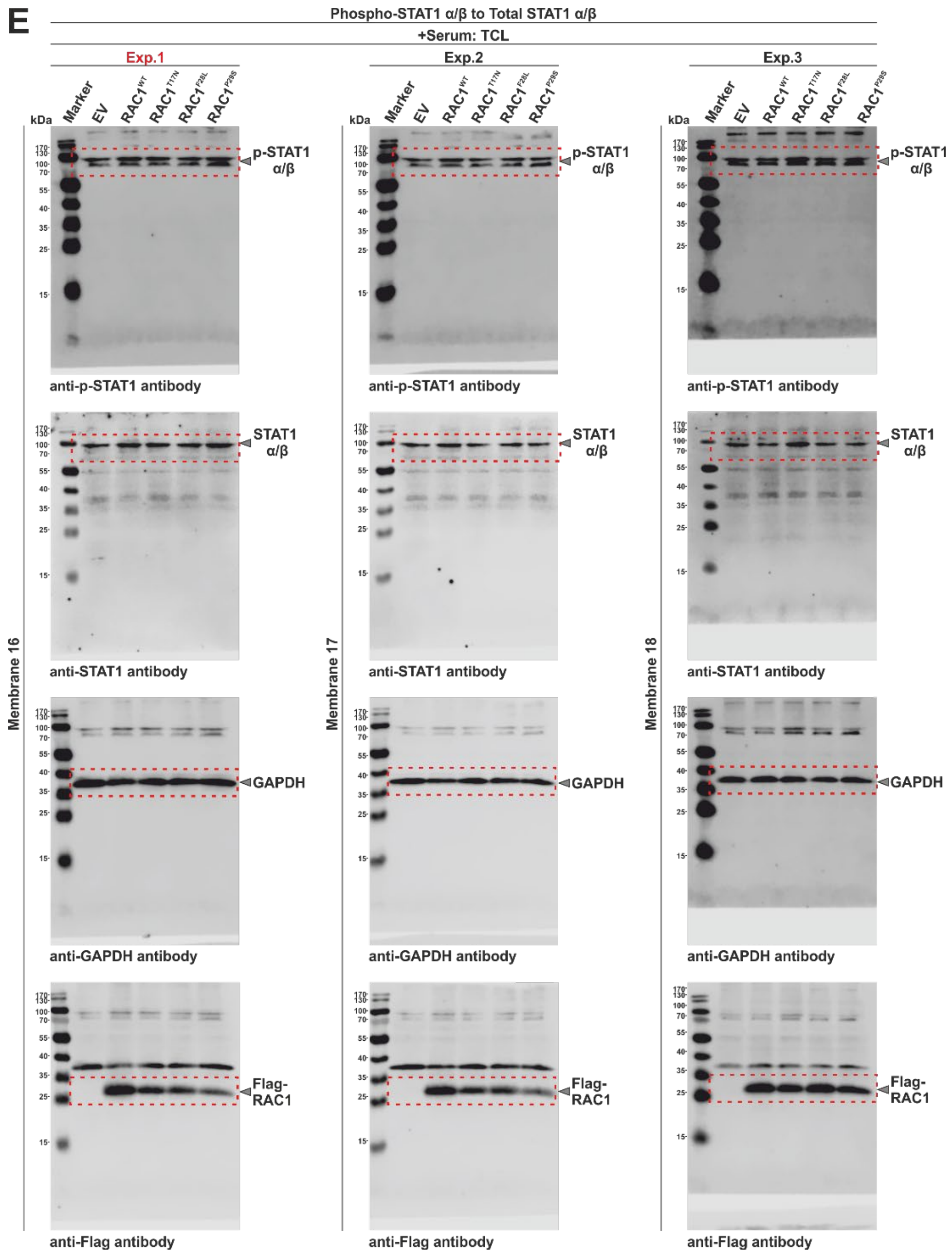


**Figure S11. Western blotting and phosphorylation analysis of ERK1/2, AKT(S473), AKT(T308), p38 MAPK, and STAT1 α/β in HEK-293T cells overexpressing RAC1 variants.** Western blot analysis was performed on serum-stimulated HEK-293T cells transiently transfected with pcDNA3.1 constructs encoding Flag-tagged RAC1 variants (WT, T17N, F28L, and P29S) alongside an empty vector (EV) control. Experiments were performed in triplicate, with each experiment visualized on nitrocellulose membranes and labeled as follows: (A) phospho-ERK1/2 to total ERK1/2 (membranes 1-3); (B) phospho-AKT (S473) to total AKT (membranes 4-6); (C) phospho-AKT (T308) to total AKT (membranes 7-9); (D) phospho-p38 MAPK to total p38 MAPK (membranes 10-15); and (E) phospho-STAT1 α/β to total STAT1 α/β (membranes 16-18). Cropped sections of the membrane shown in Figure 5 are as follows: Exp1 for AKT(T308), p38 MAPK, and STAT1 α/β; Exp2 for AKT(S473); Exp3 for ERK. Each membrane was probed sequentially, starting with the respective anti-phospho antibodies, followed by antibodies against ERK, AKT, p38 MAPK, STAT1 α/β, and GAPDH as loading controls, and the flag antibody to validate RAC1 variant expression. For p38 MAPK, separate membranes were used to detect phosphorylated and total form, as both antibodies are derived from the same host species. Detected protein bands are highlighted with dashed red boxes, and arrows indicate the corresponding molecular weight in kDa for each protein. Abbreviations:

TCL = total cell lysate; Serum+ = serum-stimulated HEK-293T cells. See the Materials and Methods section for further details.
